# TAG-IN, a modular strategy for swappable endogenous gene tagging in *Caenorhabditis elegans*

**DOI:** 10.64898/2026.01.16.700006

**Authors:** Fujia Han, Vishnu Raj, Robert D. Lemon, Qianru A. Wu, Addison Noel Stubbe, Xinyu Huang, Han Wang

**Affiliations:** Department of Biology, University of Wisconsin-Madison, Madison, Wisconsin, United States of America; Genetics Training Program, University of Wisconsin-Madison, Madison, Wisconsin, United States of America

**Keywords:** Gene tagging, RMCE, recombinase-mediated cassette exchange, CRISPR/Cas9, fusion tagging, co-expression tagging, *Caenorhabditis elegans*, bipartite expression system

## Abstract

Endogenous gene tagging allows researchers to study genes in their native genomic context, underscoring the need for efficient gene tagging approaches. Here, we report TAG-IN, a streamlined strategy for swappable gene tagging in *C. elegans*. TAG-IN begins with generating a seed allele for a gene of interest by inserting a small landing pad containing two distinct recombination sites (FRT and FRT3) into the desired tagging locus using CRISPR. Fusion or co-expression tags from reusable tag donors can then be easily swapped into the seed allele using Flp recombinase-mediated cassette exchange (RMCE). Successful TAG-IN alleles can be conveniently identified via drug selection without PCR screening. The tag-swapping step can be accomplished by simple genetic crossing, in addition to traditional microinjection. To facilitate the use of TAG-IN, we also built a library of plug-and-play tag donor strains that can be crossed with seed alleles to insert the corresponding tags and a set of vectors to construct new tag donors. The TAG-IN strategy is both modular and combinatorial: only a single seed allele for each tagging locus is needed to insert any tag; a single tag donor strain can be used to swap the tag into any compatible seed allele. Furthermore, TAG-IN allows iterative tag insertion because the FRT and FRT3 sites are preserved during RMCE, so each TAG-IN allele can serve as a landing pad for further rounds of tag exchange. Thus, TAG-IN provides an efficient, flexible, and scalable platform for endogenous gene tagging in *C. elegans,* accelerating comprehensive functional studies.

**Author summary:** Tagging genes at their native genomic locations is considered the gold standard for studying gene function, as it allows functional analysis in the original genomic context without disrupting normal regulatory processes. However, traditional CRISPR-based endogenous gene tagging methods in the model organism *Caenorhabditis elegans* are difficult to scale, especially when introducing multiple tags to the same gene, because each tag insertion requires a unique repair template delivered by microinjection, a technically challenging and labor-intensive procedure. Here, we developed TAG-IN, an efficient and modular strategy for swappable gene tagging in *C. elegans*. TAG-IN first establishes a universal landing pad at each tagging site, and then combines it with reusable tag donors to flexibly and precisely swap in desired tags via cassette exchange. TAG-IN is scalable because site-specific tag exchange can be achieved through simple genetic crosses. By making gene tagging a modular and streamlined process, TAG-IN substantially enhances the efficiency of endogenous gene tagging in *C. elegans* and provides an adaptable framework for other model organisms.

## Introduction

The genome provides instructions for proteins that carry out a wide range of molecular and cellular functions essential for life. To fully understand gene function, it is essential to determine when and where the gene is expressed, where the encoded protein goes, how the protein interacts with other partner proteins, and what happens when the protein is degraded. Genetic tools that accelerate these tasks are critical to advance our understanding of biology. In this paper, we developed a modular and efficient strategy for swappable gene tagging at endogenous loci in *Caenorhabditis elegans*, a premier model organism for biological research since the 1970s [1].

Gene tagging is a molecular genetics technique to attach specific tags to genes of interest [2]. Two major types of gene tagging, fusion tagging and co-expression tagging, serve distinct yet complementary roles in understanding the molecular and cellular mechanisms underlying gene function. Fusion tagging expresses the tag as a translational fusion with the endogenous protein, thus enabling detailed analyses of the expression, localization, interactors, and function of the tagged protein. For example, fusion tagging with GFP generates a translational reporter for directly visualizing the expression pattern and subcellular localization of the protein *in vivo* in *C. elegans* [3]. The introduction of an epitope, such as an HA tag, enables the detection of the tagged protein via immunostaining or the identification of its potential binding partners via co-immunoprecipitation [4]. Fusion tagging with TurboID, an engineered biotin ligase, can be used for proximity labeling to map protein-protein interaction networks [5,6]. Tagging with a degron, such as the AID degron or the ZF1 degron, enables degradation of the tagged protein in a spatiotemporal manner to assess its functional importance [7,8].

In contrast, co-expression tagging provides genetic access to the cells that express the target gene by producing the tag as a separate protein along with the endogenous protein. In *C. elegans*, a common strategy for co-expression involves placing the intercistronic sequence from the *gpd-2/gpd-3* operon between the target gene and the tag, which forms a synthetic bicistronic operon to achieve their simultaneous expression via SL2 *trans*-splicing [9,10]. For example, an SL2::NLS::GFP tag can be inserted into the 3’ end of the gene of interest to express nuclear-localized GFP along with the endogenous protein, thus serving as a transcriptional reporter for expression pattern analysis [11]. Co-expression tagging with transcription factors from bipartite expression systems, such as cGAL, Tet-On, Tet-Off, LexA, and the Q system [12–15], can be very powerful, as it generates a driver line that provides genetic access to the cells expressing the tagged gene. For example, a cGAL driver line can be crossed with available UAS effector lines to express the effectors in those cells for genetic rescue, expression analysis, signaling perturbation, and neuron activity manipulation [12,16]. However, such co-expression tagging with bipartite expression systems has not been implemented in *C. elegans*.

The CRISPR/Cas9 genome editing technique is a powerful tool that enables targeted modifications of the genome [17], and it has become the standard tool for endogenous gene tagging in *C. elegans* [18]. In general, CRISPR-based gene tagging in *C. elegans* involves preparing a repair template and a guide RNA for each insertion site, microinjecting the CRISPR/Cas9 genome editing mixture into the gonad, and screening for and validating successful insertions resulting from homology-directed repair following DNA cleavage by the Cas9 nuclease [18]. Over the past decade, extensive efforts have been made to enhance the efficiency of CRISPR-mediated insertion in *C. elegans* [19–26]. However, CRISPR-based gene tagging remains laborious, particularly for large tags (e.g., >1 kb) and for inserting a panel of different tags at the same locus. This is in part because each gene tagging event requires a unique repair template, which needs to be delivered to the gonad via microinjection.

In recent years, several collections of *C. elegans* strains carrying endogenously fluorescent-protein-tagged alleles of specific gene families have been established, revealing new and important insights into the functions of these gene families [26–30]. However, only 1554 protein-coding genes (about 8% of the *C. elegans* proteome) have been tagged so far [31]. Moreover, while a single tagged allele of a gene may be sufficient for an individual lab, comprehensive analysis of gene function across the *C. elegans* community often requires tagging each gene with multiple different tags for various experimental purposes. In addition, most existing tagged alleles are not designed to be easily swappable and incorporating a new tag into each locus requires an additional round of CRISPR-based insertion. Thus, there is a pressing need for efficient and flexible gene tagging strategies.

Recombinase-mediated cassette exchange (RMCE) represents a different method to insert DNA sequences into the genome using the corresponding recognition sites of the recombinase used [32]. For example, the recently developed RMCE in *C. elegans* uses the recombinase Flp and two heterospecific Flp Recognition Target sites (FRT and FRT3, 34 bp each) to efficiently generate single-copy transgenes [14,33]. Unlike the CRISPR/Cas9 genome editing technique, which requires a unique repair template consisting of the DNA insert flanked by two homology arms for each insertion locus [18], the RMCE-mediated insertion is more versatile because the same donor/repair template can be used to swap the DNA insert into different genomic loci that contain the same pair of FRT and FRT3 sites [14]. Furthermore, RMCE is generally more efficient and reliable than the CRISPR-based method for inserting large DNA sequences into the genome [14].

Here, we describe TAG-IN, a modular and scalable gene tagging strategy designed to efficiently generate various fusion and co-expression tags for virtually any protein-coding gene in *C. elegans*. With TAG-IN, only one seed allele for each tagging locus needs to be generated by CRISPR/Cas9 genome editing, and the seed allele can then serve as a landing pad that can be combined with reusable tag donors to precisely swap in any tag in parallel through RMCE, without further CRISPR editing. We demonstrated that efficient tag swapping using TAG-IN can be achieved by microinjection or simple genetic crossing, and that successful tag insertions can be conveniently identified by drug selection. We built a toolkit of reusable tag donor strains and plasmids that allow the *C. elegans* research community to readily apply the TAG-IN strategy to generate diverse tags for their genes of interest.

## Results

### Principles of a modular TAG-IN strategy for gene tagging

The general design principle of the TAG-IN strategy is to uncouple the initial modification of the tagging site from the later insertion of desired tags. First, a seed allele is generated by using CRISPR to insert a small landing pad with two distinct recombination sites (FRT and FRT3) right before the start codon or the stop codon of the target gene for N-terminal and C-terminal tagging, respectively. Second, desired tags, including fusion or co-expression tags, can then be inserted into the seed allele using Flp-mediated RMCE (Figs 1A and 1B). For fusion tagging, the resulting product is a fusion protein with the tag attached to the C-terminus or N-terminus of the endogenous protein. Co-expression tagging produces two separate protein products (the endogenous protein and the tag) in the cells that express the target gene. The TAG-IN strategy had similar but distinct designs for C-terminal or N-terminal gene tagging, as described below.

**Fig 1.**
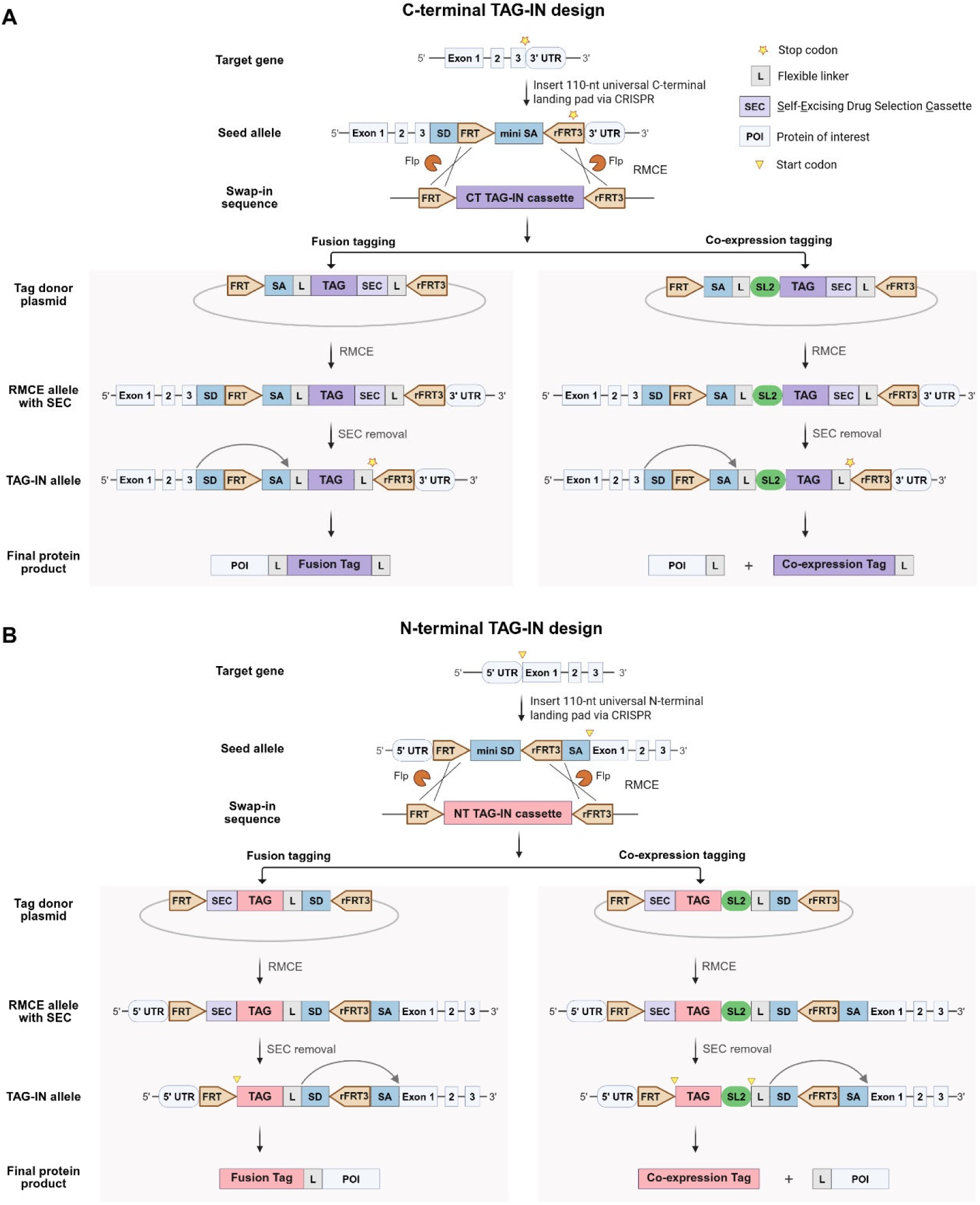
Overview of the TAG-IN strategy for both C- and N-terminal gene tagging. **(A)** Schematic of the workflow for generating a C-terminal TAG-IN allele. A C-terminal seed allele is first created by inserting a 110-nt universal C-terminal landing pad (SD-FRT-mini SA-rFRT3, refer to S1 File for the exact sequence) right before the stop codon of the target gene using CRISPR. Subsequently, different C-terminal (CT) TAG-IN cassettes flanked by FRT and rFRT3 sites can be swapped into the landing pad using Flp-mediated RMCE, generating precise C-terminal TAG-IN alleles. For C-terminal fusion tagging (lower left), each TAG-IN cassette contains a splice acceptor (SA), a flexible linker (L), a tag, a self-excision drug selection cassette (SEC), and another flexible linker (L). After RMCE and SEC removal, a C-terminal fusion TAG-IN allele will produce the endogenous protein fused with a C-terminal tag and flexible linkers (S1 Fig). Likewise, for co-expression tagging (lower right), each TAG-IN cassette consists of an SA, a flexible linker, an SL2 intercistronic sequence, a tag, a SEC, and another flexible linker. The SL2 intercistronic sequence enables simultaneous expression of the target gene and the tag from a bicistronic operon [9]. After RMCE and SEC removal, a C-terminal co-expression TAG-IN allele will produce two separate proteins, the endogenous protein and the tag, in the same cells. The 34-bp FRT site is expected to be spliced out and thus absent in the mature mRNA of each C-terminal TAG-IN allele, while the rFRT3 site will be present in the 3’UTR (S1 Fig). **(B)** Schematic of the workflow for generating an N-terminal TAG-IN allele. An N-terminal seed allele is first generated by inserting a 110-nt universal N-terminal landing pad (FRT-mini SD-rFRT3-SA, refer to S1 File for the exact sequence) right before the start codon of the target gene via CRISPR. Different N-terminal (NT) TAG-IN cassettes flanked by FRT/rFRT3 can be swapped into the seed allele via Flp-mediated RMCE. For N-terminal fusion tagging (lower left), each TAG-IN cassette contains a SEC, a tag, a flexible linker (L), and a splice donor (SD). Each N-terminal TAG-IN allele will produce the endogenous protein fused with an N-terminal tag and a flexible linker. Likewise, for N-terminal co-expression tagging (lower right), each TAG-IN cassette consists of a SEC, a tag, an SL2 intercistronic sequence, a flexible linker (L), and an SD. Each N-terminal co-expression TAG-IN allele will produce two separate proteins, the tag and the endogenous protein, in the same cells. In all N-terminal TAG-IN alleles, the rFRT3 site is expected to be removed from the mature mRNA due to splicing, while the FRT site will be present in the 5’ UTR (S2 Fig). FRT and rFRT3, Flp recognition target sequence; RMCE, recombinase-mediated cassette exchange. The curved arrow indicates splicing between the SD and SA sites. Note that the SEC is shown without its flanking lox sites for simplicity. The SEC is a temporary transgenic marker designed to minimize the labor required to identify positive knock-in worms without PCR screening [25]. The SEC is ∼5.6 kb in size and contains a hygromycin-resistance gene, a visible Rol marker, and a heat-shock inducible Cre recombinase; after self-excision, it leaves a 51-bp intron carrying a lox site (not shown here). See S1 Fig and S2 Fig for more detailed sequence information on C-terminal and N-terminal tagging, respectively.

### C-terminal TAG-IN design

For C-terminal tagging, we designed a small, universal landing pad that can be inserted immediately upstream of the stop codon of the target gene to generate a C-terminal seed allele. The landing pad is only 110 base pairs in length and includes a highly conserved splice donor in *C. elegans* [34], an FRT site, a mini splice acceptor as a spacer, and a reverse FRT3 site (Fig 1A and S1 Fig). Due to splicing and the lack of a stop codon within the spacer, each C-terminal seed allele is expected to produce the endogenous protein with a short C-terminal peptide, as it translates through a portion of the reverse FRT3 site (S1 Fig). Because of its small size, this landing pad can be easily inserted into the genome using preassembled Cas9 ribonucleoprotein (Cas9 + guide RNA) together with a synthetic single-stranded oligo repair template containing ∼35-nt long homologous arms and the 110-nt universal C-terminal landing pad [21].

We designed two types of C-terminal tag donor plasmids, distinguished by their TAG-IN cassettes, for swapping either fusion or co-expression tags into any C-terminal seed allele. Within each C-terminal tag donor plasmid, the TAG-IN cassette includes an FRT site, a splice acceptor, a flexible linker, a tag sequence for fusion tagging or an SL2::tag sequence for co-expression tagging, a transgenic marker named Self-Excising drug selection Cassette (SEC), a second flexible linker, a stop codon, and a reverse FRT3 site (Fig 1A and S1 Fig). Due to *trans*-splicing [9], the insertion of an SL2::tag will create a synthetic bicistronic operon that expresses the endogenous protein and the tag as two separate products in the same cells. The SEC, about 5.6 kb in size, contains a hygromycin resistance gene for selection, a visible roller (Rol) phenotype marker, and a heat-shock-inducible Cre recombinase. The entire SEC is flanked by two lox sites in the same orientation within a synthetic intron, allowing its self-removal by heat-shock-induced Cre expression and leaving only a 51-bp residual intron sequence [25]. The entire TAG-IN cassette in such a tag donor plasmid, including the tag and the SEC marker, can be swapped into the seed allele of the target gene via Flp-mediated RMCE between matching FRT and FRT3 sites. A final C-terminal TAG-IN allele can then be obtained after removing the SEC marker via heat shock (Fig 1A). In each final C-terminal TAG-IN allele, the FRT site is expected to be removed from the mature mRNA due to splicing, whereas the FRT3 site would remain in the 3’UTR of the tagged gene (Fig 1A and S1 Fig). Unlike original seed alleles, all final C-terminal TAG-IN alleles do not contain a short C-terminal peptide derived from a portion of the rFRT3 site, because a stop codon is included right before the rFRT site in C-terminal donor plasmids (S1 Fig).

### N-terminal TAG-IN design

The N-terminal TAG-IN strategy has the same general principle as the C-terminal tagging strategy, but uses a different landing pad for seed alleles and a distinct set of matching donor plasmids for tag swapping. We designed a 110-bp universal N-terminal landing pad that consists of an FRT site, a mini splice donor as a spacer, a reverse FRT3 site, and a highly conserved splice acceptor in *C. elegans* [34] (Fig 1B and S2 Fig). This universal N-terminal landing pad can be inserted right before the start codon of the target gene to generate a seed allele using CRISPR (Fig 1B). Due to splicing, each N-terminal seed allele would have the FRT site in the 5’UTR of the target gene (S2 Fig). We also designed two types of N-terminal tag donor plasmids for swapping either fusion or co-expression tags into any N-terminal seed allele (Fig 1B). Within each N-terminal tag donor plasmid, the TAG-IN cassette includes an FRT site, a SEC marker, a tag sequence for fusion tagging or a tag::SL2 sequence for co-expression tagging, a flexible linker, a splice donor, and a reverse FRT3 site (Fig 1B and S2 Fig). After Flp-mediated RMCE and SEC removal, a final N-terminal TAG-IN allele is generated (Fig 1B). In each N-terminal TAG-IN allele, the FRT site remains in the 5’ UTR of the target gene, whereas the FRT3 site is spliced out and thus absent in the mature mRNA (S2 Fig).

### Validation of C-terminal TAG-IN

To test the C-terminal TAG-IN strategy, we applied it to the *ceh-14* gene, which encodes a homeodomain transcription factor crucial for the development of a sleep-controlling ALA neuron in *C. elegans* [35]. We first inserted the universal C-terminal landing pad immediately before the stop codon of *ceh-14* to generate a seed allele *ceh-14(tan236)*, using a cloning-free, protein-based CRISPR/Cas9 genome editing method [36,37]. We then swapped in various tags using the original RMCE procedure [33] by microinjecting C-terminal tag donor plasmids into the gonad of *bqSi711; ceh-14(tan236)* animals to generate TAG-IN alleles. *bqSi711([mex-5p::Flp(D5)::SL2::mNG + unc-119(+)]*) is a single-copy transgene that expresses the recombinase Flp in the germline [38]. Despite a short spacer (only 12 bp) between the FRT and FRT3 sites in the seed allele, the efficiency of swapping in new tags using RMCE by microinjection was 36% (17/47, positive plates/total injection plates, two P0 each plate, S4 Fig), which is comparable to previously reported RMCE efficiency for DNA insertion into the *C. elegans* genome [14,33,39].

To test C-terminal fusion tagging, we generated a *ceh-14::gfp* TAG-IN allele and a *ceh-14::mCherry* TAG-IN allele via RMCE, essentially creating two C-terminal translational reporters for *ceh-14*. We observed that the fusion proteins CEH-14::GFP and CEH-14::mCherry localize at the nuclei of the neurons in the head and tail regions of the animal (Figs 2A and 2B), similar to the previously reported expression pattern of *ceh-14* [30].

**Fig 2.**
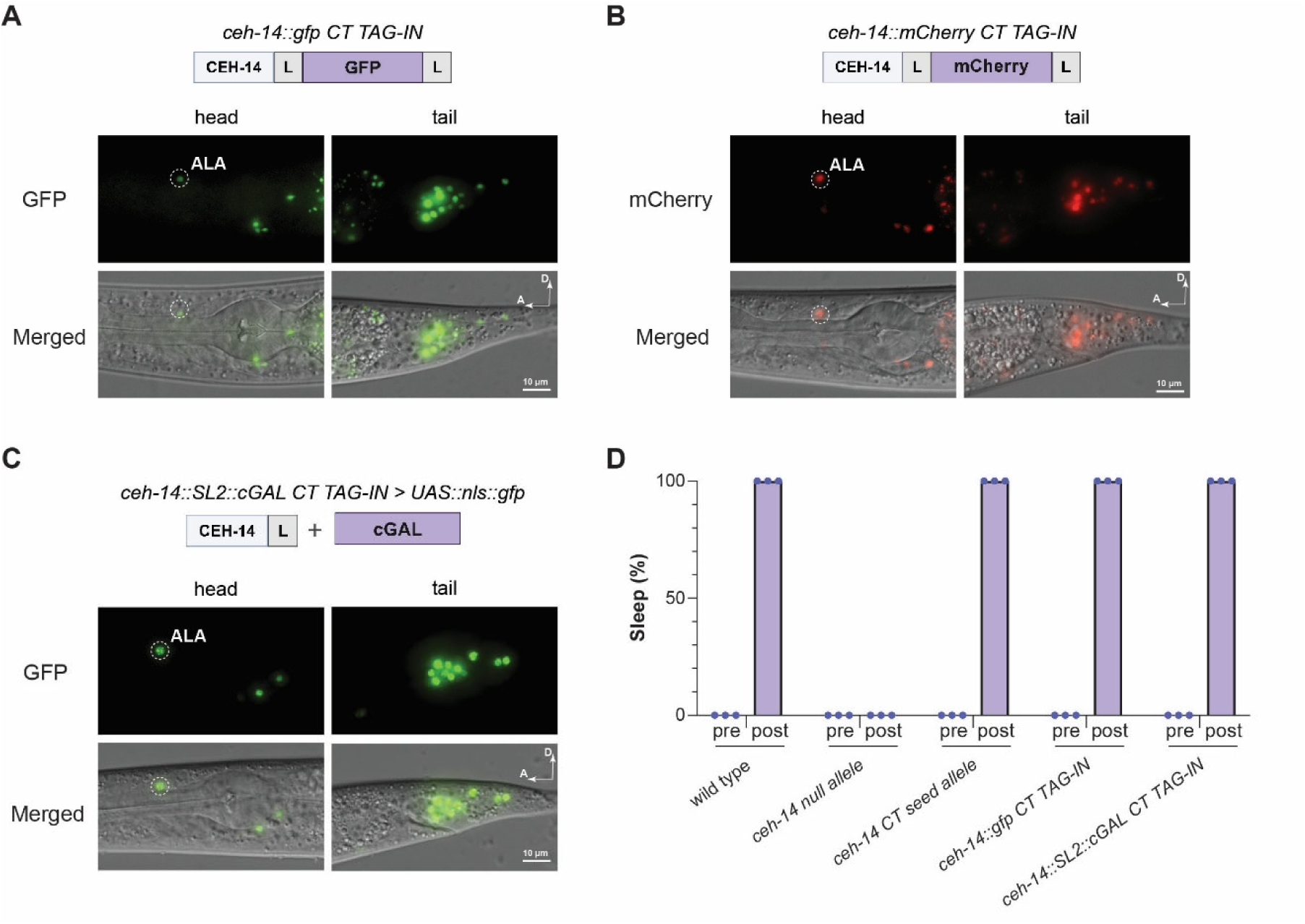
Validation of the C-terminal TAG-IN strategy in ceh-14. **(A-C)** Fluorescent and merged images of the head and tail regions of L4 animals from the following *ceh-14* C-terminal (CT) tagging strains: (A) *tan295[ceh-14::gfp CT TAG-IN]*, (B) *tan297[ceh-14::mCherry CT TAG-IN]*, and (C) *tan281[ceh-14::SL2::cGAL CT TAG-IN]*. The *ceh-14::SL2::cGAL* C-terminal TAG-IN allele was crossed with a *UAS::nls::gfp* effector (*tanSi84)*, generating a *ceh-14* transcriptional reporter. White circles indicate the location of the ALA neuron. L, flexible linker; A, anterior; D, dorsal; scale bar, 10 μm. **(D)** Quantification of pumping quiescence in adult animals with indicated genotypes and conditions. Pre, before heat shock; post, 2 hours after heat shock. All strains contain the heat-shock inducible transgene *syIs231*(*HSp::lin-3c/egf*) to trigger sleep in worms. The genotypes of worms (from left to right) are wild-type, *ceh-14*(*ot900*), *ceh-14(tan236[CT seed allele])*, *ceh-14(tan252[ceh-14::gfp CT TAG-IN ])*, and *ceh-14(tan249[ceh-14::SL2::cGAL CT TAG-IN])*. Each dot represents a group of 1-day-old adults on a single plate (N > 20 worms).

To test C-terminal co-expression tagging, we used RMCE to swap in an SL2::cGAL tag in the same C-terminal seed allele *ceh-14*(*tan236*) to generate a *ceh-14::SL2::cGAL* TAG-IN allele. Due to SL2-*trans*-splicing [9], cGAL will be expressed in the same cells where *ceh-14* is expressed, creating a cGAL driver for controlling transgene expression in those cells when combined with different UAS effectors [12]. For example, we generated a *ceh-14* transcriptional reporter by crossing this *ceh-14::SL2::cGAL* TAG-IN allele with a UAS effector line (*tanSi84[11x UAS::nls::gfp]*) to examine the expression pattern of *ceh-14*. Indeed, we observed strong NLS::GFP expression in *ceh-14*-expressing neurons in the head and tail regions (Fig 2C), similar to the expression patterns revealed by the CEH-14::GFP and CEH-14::mCherry translational reporters (Figs 2A and 2B).

Next, we tested whether C-terminal TAG-IN alleles can produce functional proteins. *ceh-14* is essential for stress-induced sleep mediated by the conserved epidermal growth factor (EGF) signaling pathway [35]. We found that, unlike the *ceh-14(ot900*) null allele, the *ceh-14* seed allele, *ceh-14::gfp* TAG-IN allele, and *ceh-14::SL2::cGAL* TAG-IN alleles all showed normal EGF-induced sleep (Fig 2D), indicating that the C-terminal seed allele and the two TAG-IN alleles do not disrupt the normal function of *ceh-14*.

### Validation of N-terminal TAG-IN

We performed similar experiments to test the N-terminal TAG-IN strategy. We inserted the 110-bp N-terminal landing pad right before the start codon of *ceh-14* to generate an N-terminal seed allele, *ceh-14*(*tan259*). For N-terminal fusion tagging, we inserted a GFP tag or an mCherry tag at the N-terminus of CEH-14 using RMCE via microinjection. In the obtained *ceh-14* N-terminal TAG-IN alleles, we observed the fusion proteins GFP::CEH-14 and mCherry::CEH-14 localized at the nuclei of the *ceh-14*-expressing neurons in the head and tail (Figs 3A and 3B). For N-terminal co-expression tagging, we used RMCE to swap *cGAL::SL2* into the N-terminal seed allele *ceh-14(tan259)*. We crossed a *cGAL::SL2::ceh-14* TAG-IN allele into the effector line *11x UAS::nls::gfp*, and observed NLS::GFP expression in *ceh-14*-expressing neurons in the head and tail regions (Fig 3C). Interestingly, the fluorescence in the ALA neuron of this reporter was hard to detect in L4 worms but could be detected in younger animals (S5 Fig). Despite being in a bicistronic operon, cGAL has a 3’ UTR derived from SL2 (the intercistronic sequence from the *gpd-2/gpd-3* operon) that is distinct from *ceh-14*’s native 3’UTR, which may explain the lower expression of cGAL in the ALA neuron at the L4 stage. Nonetheless, the *ceh-14* N-terminal seed allele, *gfp::ceh-14 TAG-IN* allele, and *cGAL::SL2::ceh-14* TAG-IN alleles all exhibited normal EGF-induced sleep (Fig 3C), indicating that the N-terminal TAG-IN strategy can also create tagged alleles that produce functional endogenous proteins.

**Fig 3.**
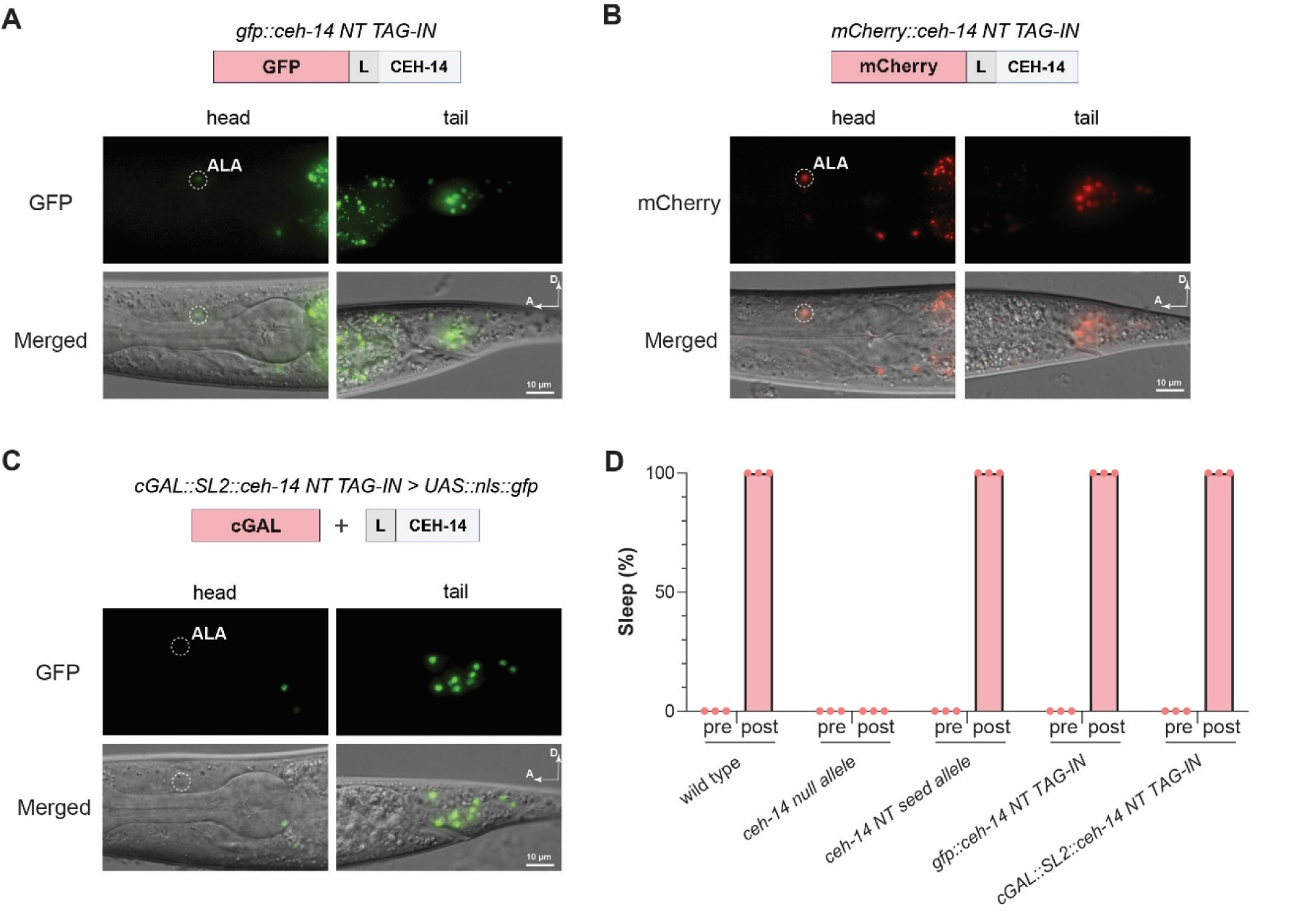
Validation of the N-terminal TAG-IN strategy in *ceh-14*. **(A-C)** Fluorescent and merged images of the head and tail regions of L4 animals from the following *ceh-14* N-terminal (NT) tagging strains: (A) *tan307[gfp::ceh-14 NT TAG-IN]*, (B) *tan309[mCherry::ceh-14 NT TAG-IN]*, and (C) *tan303[cGAL::SL2::ceh-14 NT TAG-IN]*. The *cGAL::SL2::ceh-14* N-terminal TAG-IN allele was crossed with a *UAS::nls-gfp* effector (*tanSi84)*, generating a *ceh-14* transcriptional reporter. White circles indicate the location of the ALA neuron. L, flexible linker; A, anterior; D, dorsal; scale bar, 10 μm. **(D)** Quantification of feeding quiescence in adult animals with indicated genotypes and conditions. Pre, before heat shock; post, 2 hours after heat shock. All strains contain the heat-shock inducible transgene *syIs231*(*HSp::lin-3c/egf*) to trigger sleep in worms. The genotypes of worms (from left to right) are wild-type, *ceh-14*(*ot900*), *ceh-14(tan259[NT seed allele])*, *ceh-14(tan280[gfp::ceh-14 NT TAG-IN])*, and *ceh-14(tan278[cGAL::SL2::ceh-14 NT TAG-IN])*. Each dot represents a group of 1-day-old adults on a single plate (N > 20 worms).

Taken together, these results demonstrate that the TAG-IN strategy can efficiently insert different fusion or co-expression tags into a C-terminal or N-terminal seed allele of the target gene via RMCE and the resulting TAG-IN alleles can retain normal function of the target gene.

### RMCE-mediated TAG-IN via genetic crossing

We have recently demonstrated that RMCE-mediated insertion into the *C. elegans* genome can also be achieved by genetic crossing [39]. To lower the energy barriers for generating endogenous gene tagging alleles using the TAG-IN strategy, we sought to determine whether a similar genetic crossing approach could bypass the labor-intensive and technically challenging step of microinjecting individual tag donor plasmids into each seed allele for RMCE to occur. Because each TAG-IN donor plasmid is reusable, and plasmids injected into the gonad of adult hermaphrodites can be assembled into large transmissible extrachromosomal arrays [40], we tested whether each donor plasmid could be injected only once to establish a reusable tag donor array that could then be crossed with strains carrying a seed allele and a separate germline-expressing Flp transgene to insert the tag via RMCE.

To this end, we designed tag donor arrays in which each array contains a tag donor plasmid and two histamine-based negative selection markers. The SEC marker in each tag donor plasmid causes transgenic animals carrying either the extrachromosomal arrays or successful insertion of the tag cassette to display a Rol phenotype and hygromycin resistance [25]. In the presence of histamine, the two negative selection markers kill worms carrying the arrays by causing feeding cessation and uncoordinated movement via inhibition of both the nervous system and the pharyngeal muscle [39,41]. The combination of both positive and negative selection markers in donor arrays was designed to provide a convenient way to isolate TAG-IN alleles from genetic crosses, as described below.

An overview of the procedure for generating TAG-IN alleles through RMCE via genetic crossing is shown in Fig 4A. On day 0, we cross males that carry both a seed allele and a germline-expressing Flp transgene with hermaphrodites carrying a tag donor array to generate the cross progeny that contain the seed allele, the germline-expressing Flp transgene, and the tag donor array. RMCE is likely to occur in the germline of these cross progeny: the Flp recombinase first resolves the array into small circular DNA molecules and then swaps the TAG-IN cassette into the seed allele using the matching FRT and FRT3 sites [14]. On day 3, we add hygromycin to positively select for such cross progeny, as well as other worms carrying the tag donor array. On day 6, we apply histamine to negatively select against animals carrying the donor array. On day 9, we single out surviving Rol animals showing robust pumping into plates without drug selection. In the next generation, Rol animals are then singled out again from putative positive plates containing ∼75% Rol progeny to establish homozygous Rol lines. After SEC removal via heat shock, we get the TAG-IN alleles with the tag inserted.

**Fig 4.**
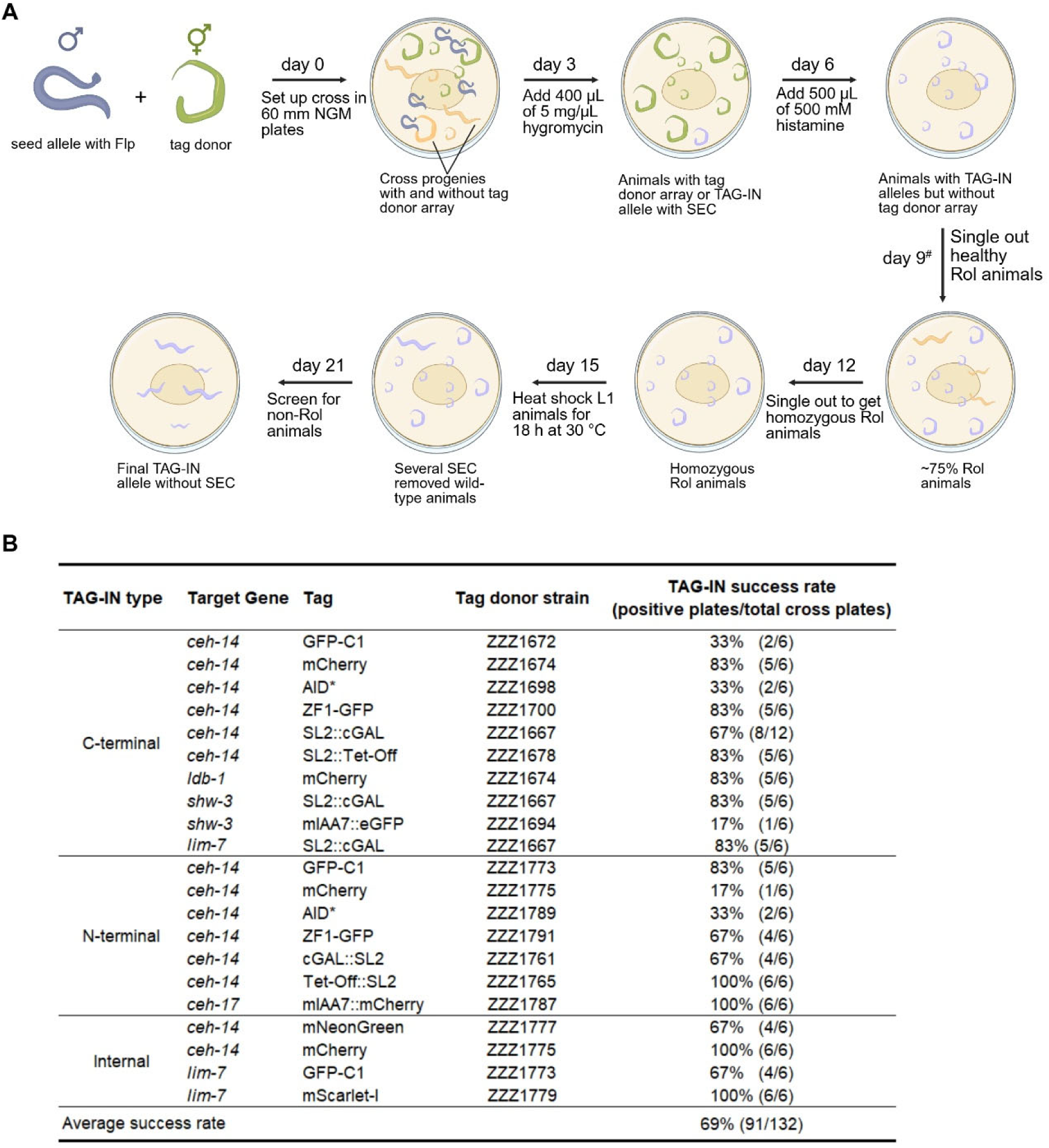
Generation of TAG-IN alleles through RMCE via genetic crossing. **(A)** Schematic workflow for generating a new TAG-IN allele by crossing a seed allele with a compatible tag donor array. On day 0, crossing males carrying both the seed allele and a germline-expressing Flp transgene (either double heterozygous or double homozygous) with hermaphrodites carrying the tag donor array (with a dominant Rol marker) in half a dozen 60 mm NGM plates, each containing 10 mL of media. Tag insertion via RMCE starts to occur in the germline of the cross progeny on day 3, where the Flp recombinase resolves the donor array into individual small TAG-IN donor DNAs and then swaps the tag into the landing pad [14]. The sequential dual drug selection with hygromycin followed by histamine in the following days, as depicted, effectively selects for progeny with successful TAG-IN cassette insertion while lacking the tag donor arrays. Typically, nine days after the cross, 1-4 candidate TAG-IN worms per positive plate that show the Rol and robust pumping phenotypes are singled out into individual NGM plates for further confirmation in the next generation. #, in our experience, surviving Rol animals may show up 6-14 days after the cross, depending on when the RMCE event occurs. Positive worms with a successful TAG-IN insertion are expected to produce ∼75% Rol progeny in the next generation. Progeny from these plates are then singled out again to obtain homozygous Rol animals. After SEC removal via heat shock, wild-type (non-Rol) animals are recovered to isolate final TAG-IN alleles. TAG-IN alleles isolated from the same cross plate are considered to be the same line. **(B)** Table showing the success rates of generating TAG-IN alleles from the genetic crosses between indicated seed alleles and tag donor strains. *bqSi711* is the germline-expressing Flp transgene used for RMCE.

To test our crossing procedure to generate new TAG-IN alleles, we applied it to a C-terminal *ceh-14* seed allele and an N-terminal *ceh-14* seed allele. We observed high efficiency for both C-terminal and N-terminal gene tag swapping by RMCE via genetic crossing, with an average success rate of 64% (27/42, positive plates/total cross plates) and 61% (22/36), respectively (Fig 4B and S1 Table), which were comparable, if not higher, to the efficiency of RMCE via direct microinjection.

### Robustness of TAG-IN performance across multiple gene loci

To further assess whether the TAG strategy is generally applicable for gene tagging at other endogenous loci, we applied it to four additional genes (*lim-7*, *ceh-17*, *shw-3*, and *ldb-1*). We found that the average success rate of generating the corresponding TAG-IN alleles of these genes via RMCE genetic crossing was 73% (22/30), comparable to that of *ceh-14* (63%, 49/78). Among all tags and genes examined, the average success rate of TAG-IN was 69% (Fig 4B and S1 Table), demonstrating the robustness of the TAG-IN strategy.

To assess the fidelity of tag insertion using TAG-IN, we used PCR and sequencing to genotype 40 independent TAG-IN alleles generated by RMCE via genetic crossing, representing 21 different tagged alleles across eight tagging sites. We found that 38 out of 40 TAG-IN alleles (95%) had precise insertion of the expected tag sequences (S1 Table). Genotyping PCR for the remaining two TAG-IN alleles failed (S1 Table). Similar rare unexpected insertions have also been observed in a previous study on RMCE-mediated single-copy transgene insertion, which may be due to incomplete RMCE or integration of a much larger fragment formed during donor array formation [33]. Taken together, these results demonstrate that the TAG-IN strategy allows efficient and precise tag insertions at multiple genes through RMCE by genetic crossing and indicate that it can be used to generate tagged alleles for other protein-coding genes in the *C. elegans* genome.

### Iterative tag swapping using TAG-IN alleles as landing pads

Since each TAG-IN allele is flanked by the same FRT and FRT3 sites as the original seed allele, we tested whether TAG-IN alleles can also serve as landing pads for tag swapping through additional rounds of RMCE. To do so, we used the *ceh-14(tan307[gfp::ceh-14 NT TAG-IN])* allele as a test case. Using RMCE via genetic crossing, the success rates of replacing GFP in the *ceh-14(tan307)* TAG-IN allele with either mScarlet-I or mCherry were 58% (7/12) and 50% (6/12), respectively (S6B Fig). Thus, the TAG-IN strategy enables iterative RMCE to replace the existing tag in a TAG-IN allele with a new one via simple genetic crossing.

### A new transgene line for germline-expressing Flp recombinase

To conveniently generate TAG-IN alleles for genes across all six chromosomes in *C. elegans* through RMCE via genetic crossing as described above, Flp-expressing transgenes on different chromosomes are needed. Initially, we used *bqSi711 ([mex-5p::Flp(D5)::SL2::mNG::glh-2 3’UTR + unc-119(+)]*), a single-copy Flp-expressing transgene inserted at the *cxTi10882* locus on Chr IV [38], which worked robustly for generating TAG-IN alleles at multiple loci (Fig 4B). However, due to genetic linkage, *bqSi711* is not suited for generating TAG-IN alleles for those genes on Chr IV via genetic crossing. To address this, we generated an alternative single-copy transgene, *tanSi334,* inserted at the *ttTi5605* locus on Chr II using MosSCI [42]. *tanSi334* expresses Flp robustly in the germline, as indicated by the green fluorescence from mNeonGreen (mNG) (S6A Fig). The performance of *tanSi334* in generating TAG-IN alleles through RMCE via genetic crossing was comparable to that of *bqSi711* (S6B Fig).

### A TAG-IN toolkit of donor plasmids and strains

To facilitate the adoption and increase the accessibility of the TAG-IN strategy, we built 42 tag donor plasmids (Table 1, S3 Table, and S1 File): a collection of 30 fusion tag donor plasmids (15 for C-terminal tagging and 15 for N-terminal tagging) for inserting various commonly used tags, including fluorescent proteins, epitope tags, degrons, and enzymatic tags; a library of ten co-expression tag donor plasmids (six for C-terminal tagging and six for N-terminal tagging) for inserting the transcription factors from six bipartite expression systems—cGAL, Tet-On, Tet-Off, LexA, QF, and QF2—all of which have been shown to work in *C. elegans* [12–15]. Furthermore, we injected each of these tag donor plasmids to establish 42 donor extrachromosomal arrays (Table 1 and S2 Table), which can be directly used to cross with compatible C-terminal or N-terminal seed alleles to swap in the corresponding tags through RMCE.

**Table 1.**
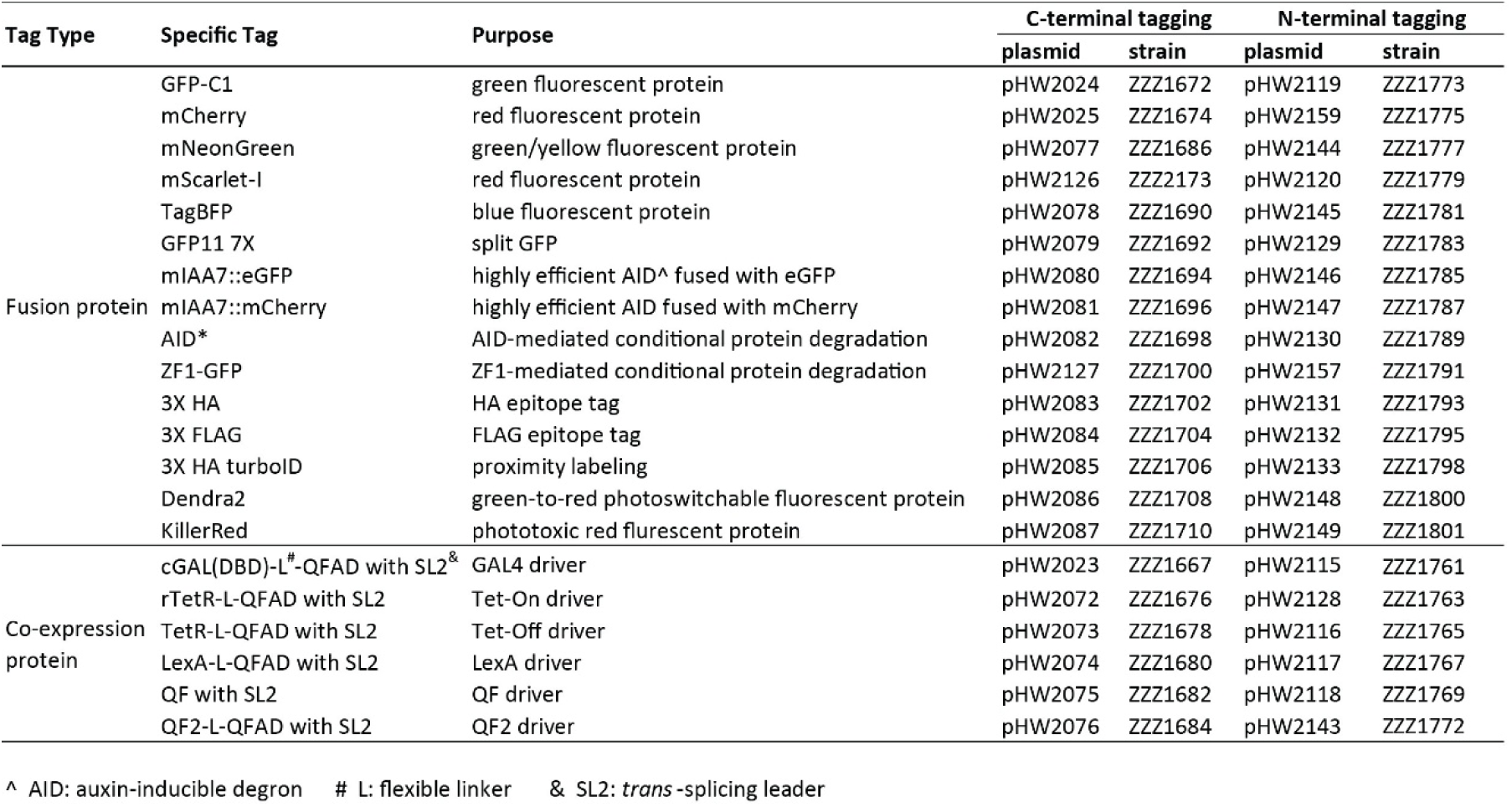
Tag donor plasmids and tag donor strains generated in this study.

### A set of TAG-IN destination vectors for cloning new tags

To facilitate future molecular cloning to build new TAG-IN donor plasmids, we designed four major types of destination vectors: one for C-terminal fusion tagging, one for C-terminal co-expression tagging, one for N-terminal fusion tagging, and one for N-terminal co-expression tagging (Figs 5A and 5B). For each vector type, we created three different versions, with each one containing one of the three available SEC (flanked by a pair of loxP, lox2272, or lox511I sites). For constructing new tag donor plasmids, we followed the previously established convention for SEC selection: the loxP version for green tags, the lox2272 version for red tags, and the lox511I version for all other tags [25].

**Fig 5.**
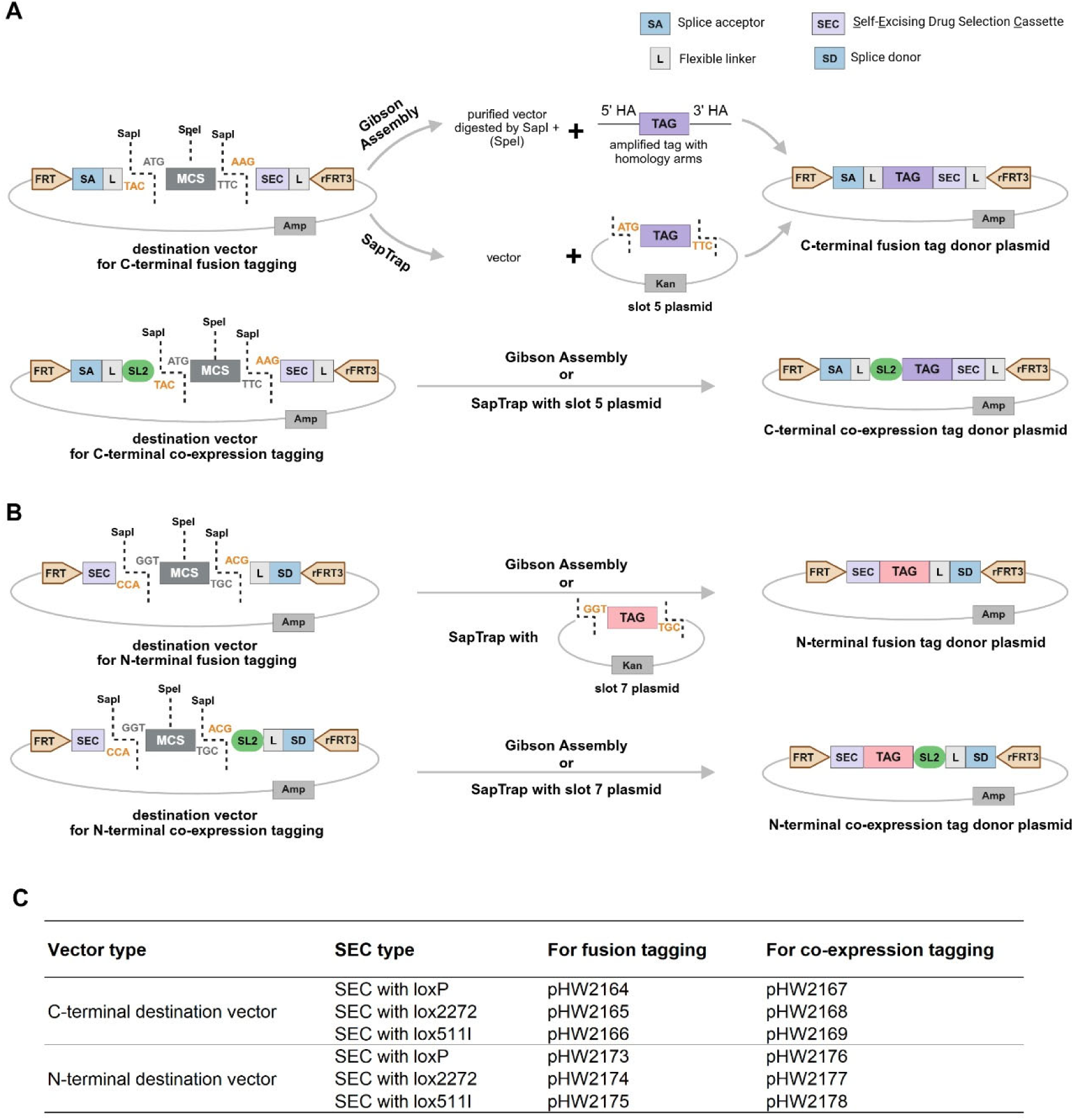
TAG-IN destination vectors to construct new tag donor plasmids. **(A and B)** Scheme for constructing new tag donor plasmids using TAG-IN destination vectors created in this study. (A) Construction of new C-terminal fusion or co-expression tag donor plasmids using the corresponding destination vectors. A destination vector is linearized with the restriction enzyme SapI (optionally with another enzyme, SpeI), purified, and then assembled with a tag fragment containing appropriate homology arms using Gibson assembly. Alternatively, a destination vector and a SapTrap slot 5 plasmid (containing a tag, flanked by the adaptor sequences ATG and AAG) can be directly combined to construct a new tag donor plasmid through SapTrap. (B) Construction of new N-terminal fusion or co-expression tag donor plasmids using the corresponding destination vectors. Cloning strategies are the same as those described in (A). The difference is that, instead of a slot 5 plasmid, a slot 7 plasmid (containing the tag, flanked by the adaptor sequences GGT and ACG) is used for SapTrap. **(C)** Table listing 12 TAG-IN destination vectors for constructing new tag donor plasmids. We have built four types of vectors for fusion tagging or co-expression tagging at the C-terminus or the N-terminus. Each vector type has three variants, each of which contains a SEC marker flanked by one of three lox sites: loxP, lox2272, or lox511I.

To construct a new tag donor plasmid, simply clone a single DNA fragment encoding the tag with a start codon but lacking a stop codon into the corresponding destination vector using Gibson Assembly (Figs 5A and 5B). Alternatively, new tags can also be conveniently inserted into these vectors using SapTrap [43], a Golden Gate Assembly strategy (Figs 5A and 5B). Existing tags in the SapTrap slot 5 entry vector (historically the FP slot, flanked by the adaptor sequences ATG-AAG [44]) can be directly cloned into either the C-terminal fusion or the co-expression vectors; existing tags in the SapTrap slot 7 entry vector (historically the NT tag linker slot, flanked by the adaptor sequences GGT-ACG [44]) can be directly cloned into either the N-terminal fusion or co-expression tagging vectors.

### Repurposing N-terminal TAG-IN donors for internal tagging

For various reasons, a small fraction of genes may not tolerate being tagged at either the C-terminus or N-terminus to generate functional fusions. A recent systematic survey of currently available endogenous gene tagging in *C. elegans* reveals that 2.9% are internal tagging [31]. We sought to adapt the N-terminal TAG-IN for internal tagging by introducing a small but critical modification to the landing pad sequence for internal seed alleles (Fig 6A). Specifically, we designed a 112-bp universal internal landing pad, essentially the 110-bp universal N-terminal landing pad plus two guanines (GG) at its 5’ end (S3 Fig and S1 File). This 112-bp cassette must be inserted in-frame into an exon of the target gene to generate an internal seed allele, which is fully compatible with N-terminal TAG-IN fusion donors (Fig 6A). The two added guanines (GG), together with the downstream 34-bp FRT site as well as the AAG and GGT adaptors from SapTrap cloning of the SEC marker, encode a 14-amino-acid protein linker (“GKFLFSRKYRNFKG”) immediately following the insertion site, preserving the reading frame of the internal fusion protein (S3 Fig).

**Fig 6.**
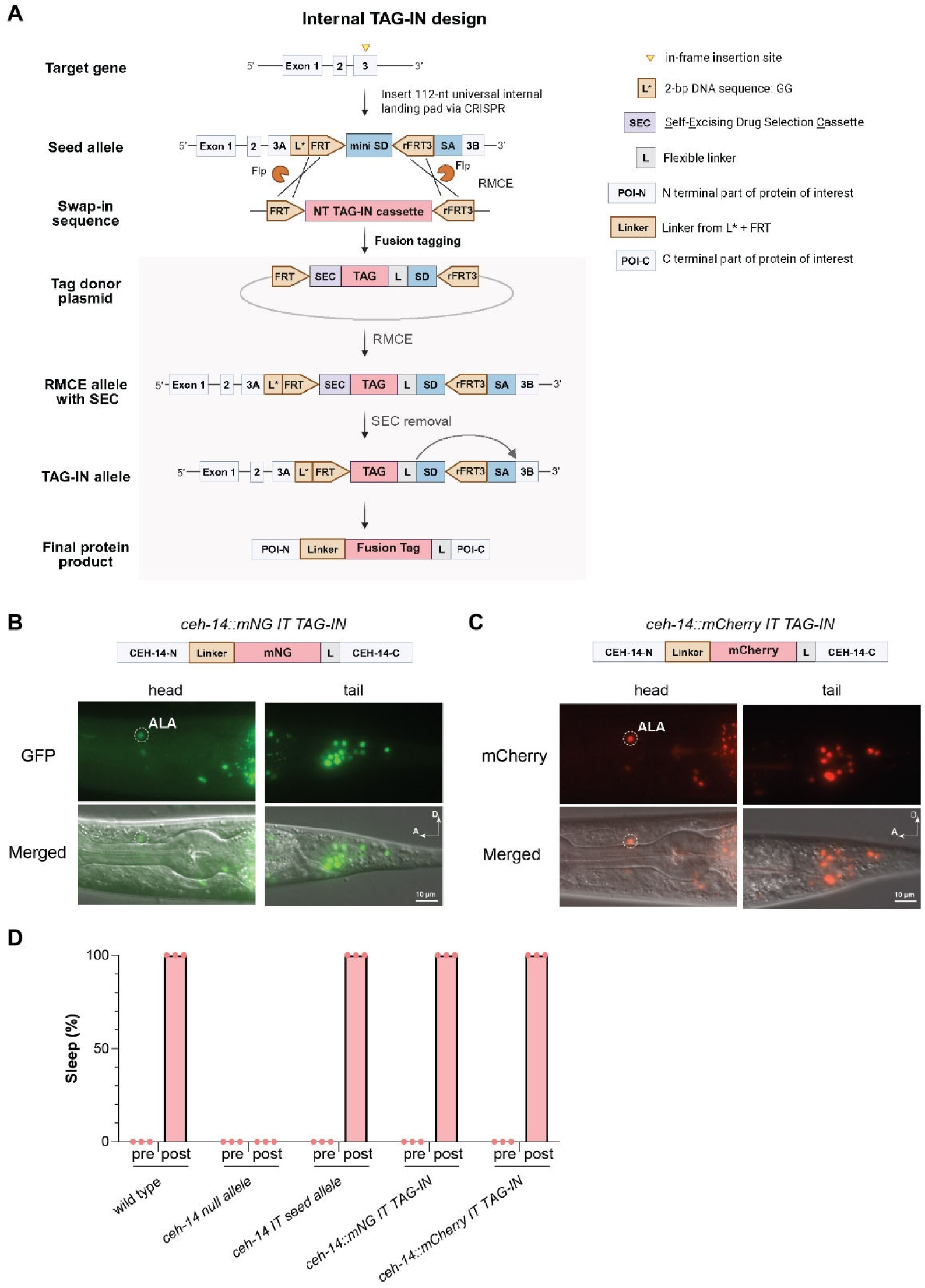
Validation of the internal TAG-IN strategy in *ceh-14*. **(A)** Schematic of the workflow for generating an internal TAG-IN allele. An internal seed allele is first generated by inserting the 112-nt universal internal landing pad (L*-FRT-mini SD-rFRT3-SA, refer to S1 File for the exact sequence) in-frame into an exon of the target gene via CRISPR. Such internal seed alleles are fully compatible with N-terminal TAG-IN donors for fusion tagging: N-terminal TAG-IN cassettes flanked by FRT and rFRT3 sites can be swapped into any internal seed allele to replace the mini SD via Flp-mediated RMCE. Each internal fusion TAG-IN allele will produce the endogenous protein fused to an internal tag with flanking linkers. In both seed alleles and internal TAG-IN alleles, the “L*-FRT” sequence will produce a 12-amino-acid peptide linker (“GKFLFSRKYRNF”) immediately following the tagging site. FRT and rFRT3, Flp recognition target sequence; RMCE, recombinase-mediated cassette exchange. Note that the SEC is shown without its flanking lox sites for simplicity. After self-excision, the SEC leaves a 51-bp intron carrying a lox site (not shown here). See S3 Fig for more detailed sequence information on internal TAG-IN. **(B-C)** Fluorescent and merged images of the head and tail regions of L4 animals from the following *ceh-14* internal (IT) tagging strains: (B) *tan384[ceh-14::mNG IT TAG-IN]*, (C) *tan387[ceh-14::mCherry IT TAG-IN]*. White circles indicate the location of the ALA neuron. L, flexible linker; A, anterior; D, dorsal; scale bar, 10 μm. **(D)** Quantification of feeding quiescence in adult animals with indicated genotypes and conditions. Pre, before heat shock; post, 2 hours after heat shock. All strains contain the heat-shock inducible transgene *syIs231*(*HSp::lin-3c/egf*) to trigger sleep in worms. The genotypes of worms (from left to right) are wild-type, *ceh-14*(*ot900*), *ceh-14(tan367[IT seed allele])*, *ceh-14(tan384[ceh-14::mNG IT TAG-IN])*, and *ceh-14(tan387[ceh-14::mCherry IT TAG-IN])*. Each dot represents a group of 1-day-old adults on a single plate (N > 20 worms).

We first tested the internal TAG-IN strategy in *ceh-14*. We used CRISPR to generate an internal seed allele, *ceh-14(tan367)*. This allele was subsequently used as a landing pad and combined with existing N-terminal tag donor arrays (Fig 4B and Table 1) to swap in either mNG or mCherry via RMCE to generate two internal TAG-IN alleles, *ceh-14(tan384[ceh-14::mNG IT TAG-IN])* and *ceh-14(tan387[ceh-14::mCherry IT TAG-IN])*, respectively. We found that these two *ceh-14* internal TAG-IN alleles were functional and that the corresponding fusion proteins localized at the nuclei of *ceh-14*-expressing neurons in the head and tail regions (Figs 6B-6D). We tested the internal TAG-IN strategy in another gene, *lim-7,* and found that it also worked robustly (Fig 4B). Taken together, the TAG-IN strategy can also be used for internal fusion tagging.

## Discussion

Here, we present TAG-IN, a modular and versatile strategy for swappable endogenous gene tagging in *C. elegans*. In TAG-IN, each tagging locus is edited only once by CRISPR to insert a small RMCE landing pad, generating a seed allele. Desired fusion or co-expression tags can then be repeatedly swapped into the locus to generate TAG-IN alleles via RMCE by combining the seed allele with reusable tag donors. We established a convenient pipeline for streamlined RMCE-mediated tag swapping using simple genetic crossing coupled with dual drug selection. We showed that TAG-IN alleles support iterative RMCE to replace existing tags with new ones. Finally, we constructed a ready-to-use library of 42 reusable tag donors and a toolkit of destination vectors for generating new tag donor plasmids. Because of its modular design, combinatorial features, and the ease of RMCE via genetic crossing, TAG-IN provides a scalable platform for efficient gene tagging at endogenous loci in *C. elegans*.

While both injection and crossing are efficient options for the tag swapping step in the TAG-IN procedure, crossing is generally preferred in practice for most labs because it is simpler to implement, particularly for tags with available donor arrays and for new tags that don’t yet have donor arrays but need to be inserted at multiple genomic loci. Injection provides a useful alternative when crossing is not feasible or when the donor array cannot be established due to toxicity, although it requires construction of a parental strain homozygous for both the seed allele and a germline-expressing Flp transgene.

Our results demonstrate that a 12-bp spacer between the FRT and FRT3 sites of the landing pad in seed alleles is sufficient to support efficient RMCE to insert tags of different sizes. Beyond gene tagging, the general design principle of the TAG-IN strategy can potentially be adapted to insert long DNA sequences into defined loci in the *C. elegans* genome, since short sequences are generally easier to insert than long sequences via CRISPR-mediated homology-directed repair [18,21]. For example, a small landing pad flanked by FRT and FRT3 sites can be first inserted into an intron of a target gene, and the resulting allele can be used for RMCE to swap in previously reported large gene trap cassettes [39] to generate new gene trap lines. Likewise, a similar cassette can also be introduced into well-characterized genomic loci known to support reliable transgene expression [42,45] to generate additional landing pads for RMCE-mediated insertion of single-copy transgenes.

### Advantages of the TAG-IN strategy

First, TAG-IN is modular and scalable. Only one seed allele needs to be created for each tagging locus, and each tag donor plasmid/strain only needs to be constructed once. Seed alleles and tag donors can then be mixed and matched to insert the tag through RMCE. Our design of combining simple genetic crossing and drug selection to obtain TAG-IN alleles makes endogenous gene tagging in *C. elegans* easier and allows multiple tags at the same locus, or the same tag across different loci, to be generated in parallel, especially once seed alleles and tag donor strains have been established. TAG-IN is particularly useful when multiple tags for a given locus are needed, as is often the case in the *C. elegans* community where different labs may require different tags at the same locus. Second, TAG-IN is efficient for various insertion sizes. The high efficiency of TAG-IN observed for inserting diverse tags (0.3kb–1.6 kb after removing the 5.6 kb SEC marker) in this study is consistent with previous work showing robust performance of RMCE-mediated single-copy transgene insertion into the *C. elegans* genome, including large inserts ranging from 9.3 kb to 13.9 kb [14,33]. Third, TAG-IN is precise because it uses cassette exchange to swap in tags from donor constructs. Fourth, TAG-IN is iterative and future-proof. Because the flanking FRT and FRT3 sites are preserved during RMCE, each TAG-IN allele becomes a new seed allele, serving as a landing pad for repeated rounds of RMCE-mediated tag replacement. As improved or new tags are developed, existing TAG-IN alleles can be readily converted into updated TAG-IN alleles with these new tags via RMCE, without additional CRISPR genome editing. Fifth, the TAG-IN strategy is versatile because the same seed allele can be used for both fusion and co-expression tagging. In particular, co-expression tagging can convert the seed allele into drivers of bipartite expression systems, which can be combined with appropriate effector lines to genetically label or manipulate cells where the tagged gene is expressed. Sixth, TAG-IN alleles can be further modified for functional studies. For example, insertion of an FRT site, a lox site, or a reverse FRT3 site in an early intron of a C-terminal TAG-IN allele or in the 3’ UTR of an N-terminal TAG-IN allele can create a conditional knockout allele of the target gene, which can be crossed with existing tissue-specific Flp or Cre lines for tissue-specific knockout experiments [46,47].

### Potential limitations of the TAG-IN strategy

First, as synthetic single-stranded oligos longer than 200 nucleotides are costly and usually of lower purity, if there are no ‘NGG’ PAM sites near the start codon or the stop codon of the target gene (e.g., > 30 bp away), it can be challenging to use such oligos as repair templates to create seed alleles. In this scenario, one potential solution is to use engineered Cas9 variants or other CRISPR nucleases, such as Cpf1/Cas12a, that recognize other PAM sites located closer to the start codon or the stop codon [48–50]. Alternatively, *in vivo* template switching using multiple single-stranded oligos [51], custom-made long single-stranded DNA templates [23], or synthetic double-stranded DNA templates can be used for CRISPR/Cas9-mediated insertion of small landing pads to generate seed alleles. Second, due to a transcriptional terminator at the beginning of the SEC marker, its insertion in N-terminal and internal TAG-IN alleles will generate a temporary loss-of-function allele of the target gene [25]. Homozygotes of such TAG-IN alleles before SEC removal cannot be recovered if loss-of-function alleles of the target gene are lethal. In such cases, multiple strategies, such as the use of balancers, can be employed to track the TAG-IN alleles during SEC removal. Another option is to replace the SEC marker in tag donor plasmids with a recently developed Nested, Self-Excising selection Cassette (NSEC) to alleviate this limitation [52]. Third, although our results demonstrated that TAG-IN can generate functional tagged alleles, it is worth noting that, as with any tagging approach, introducing a tag to an endogenous gene may perturb its normal function. The extent of such issues may depend on the specific tag and/or the insertion site. Therefore, it is critical to carefully select both the tag and insertion and to confirm that TAG-IN alleles retain normal function before using them for downstream functional studies. To generate new seed alleles and TAG-IN alleles, it is recommended to follow the best practices for endogenous gene tagging in *C. elegans*, as recently reviewed [31].

### Comparison of TAG-IN with other gene tagging methods

The CRISPR-based gene tagging can be faster, as the desired tagged alleles are obtained directly in the knock-in strains [18]. Tag swapping through CRISPR is feasible but requires repeated microinjection [28]. The TAG-IN strategy may take longer to get the first TAG-IN alleles for a gene due to the two-step process. However, once a seed allele has been established, TAG-IN is easier to implement and more efficient and versatile than CRISPR-based tag insertion, because the same seed allele can be combined with different TAG-IN donors via simple genetic crossing to generate multiple tagged alleles in parallel.

A recent preprint describes a related but orthogonal approach, PhIT, which uses both PhiC31 integrase and Dre recombinase to insert a tag carrying an *attP* site into the target gene that has been previously edited to contain an *attB* site [53]. Similar to TAG-IN, PhIT allows insertion of multiple tags in parallel at a given locus by repeatedly using the same original landing pad carrying the *attB* site. However, PhIT and TAG-IN are different in several aspects. First, because it relies on the *unc-119(+)* selection marker, PhIT needs to be performed in an *unc-119* mutant background; TAG-IN can be applied in virtually any genetic background. Second, PhIT leaves peptide scars translated from the resulting *attL* and *attR* sequences flanking the inserted tag; due to splicing, TAG-IN does not leave peptide scars from the FRT and FRT3 sites for C-terminal and N-terminal tagging, respectively. Third, unlike TAG-IN, PhIT does not support iterative tag editing of tagged alleles because the *attB* site within the original landing pad is disrupted following each tag insertion.

### Outlook of the TAG-IN strategy

Despite the efforts of the community over the past decade, only 1554 genes, which represent about 8% of the *C. elegans* protein-coding genes, have been endogenously tagged [31], partly due to the lack of an efficient and high-throughput method for endogenous gene tagging. Our TAG-IN strategy is plug-and-play, swappable, precise, and scalable. We envision that with a genome-wide collection of seed alleles, our TAG-IN strategy can effectively tag the entire *C. elegans* proteome with diverse tags using the currently available as well as future new tag donor strains. Thus, the TAG-IN strategy adds a powerful gene-tagging tool to the *C. elegans* genetic toolkit and will facilitate functional studies of gene expression and function.

## Materials and methods

### *C. elegans* strain maintenance

All *C*. *elegans* strains were grown and maintained at room temperature on Petri plates as described [54], unless otherwise specified in the text. Each 60 mm Petri dish containing 10 mL of NGM agar was seeded in the center with an overnight culture of the *E*. *coli* strain OP50 grown in LB media. A complete list of strains used and generated in this study is listed in Supplementary S2 Table.

### Molecular biology

Oligos and double-stranded DNA fragments used in this study were synthesized (Integrated DNA Technologies, Coralville, IA, or Twist Bioscience, San Francisco, CA). Plasmid cloning was done via Gibson assembly using the NEBuilder HiFi DNA Assembly Master Mix (New England Biolabs, Ipswich, MA), or via a Golden Gate cloning method called SapTrap [43]. For SapTrap, LguI (Thermo Scientific, Waltham, MA) or BspQI-HF (New England Biolabs, Ipswich, MA), two isoschizomers for the restriction enzyme SapI, were used instead. Transformation was mostly done using home-made DH5α competent cells. Positive clones were verified by Oxford Nanopore long-read sequencing (Plasmidsaurus, Eugene, OR). Plasmids and oligos used and generated in this study are listed in S3 Table and S4 Table, respectively. Annotated DNA sequences of all TAG-IN donor plasmids and destination vectors are included in S1 File.

### Generation of transgenic animals

Injection solutions with various components were injected into the gonad of adult hermaphrodites using a standard microinjection procedure [40] to create transgenic animals with different strategies described below.

### Generation of seed alleles

The seed alleles in this study were generated through a cloning-free, protein-based CRISPR/Cas9 genome editing protocol with the *dpy-10* co-conversion marker [36,37]. Briefly, Individual components, including Cas9 protein (QB3 MacroLab, UC Berkeley), tracrRNA (Alt-R™ CRISPR-Cas9 tracrRNA, 20 nmol, IDT), gRNA (Alt-R™ CRISPR-Cas9 crRNA, 2 nmol, IDT), and ssDNA repair templates (Ultramer DNA Oligo, 4 nmol, IDT) were purchased for CRISPR. A typical injection mixture contains 2 μM of Cas9 protein, 0.5 μM of crRNA for *dpy-10*, 2.0 μM of crRNA for the target gene, 2.5 μM of tracrRNA, 0.5 μM of the repair template for *dpy-10*, and 0.5 μM of the repair template for the target gene, brought to a total volume of 20 μL with nuclease-free water (NEB). For most experiments, we injected the CRISPR mixture into ∼20 N2 worms. Putative seed alleles were identified by PCR genotyping F_2_ progenies of individually F_1_ Rol animals singled out from jackpot plates, followed by gel electrophoresis. PCR products with predicted insertion sizes were purified and sent for Sanger sequencing (Functional Biosciences, Madison, WI) to identify the correct clone with the seed allele. Confirmed lines were singled out again to obtain non-Rol animals that are homozygous for the seed allele in the next generations. In general, the Custom Alt-R™ CRISPR-Cas9 guide RNA design tool from IDT (https://www.idtdna.com/site/order/designtool/index/CRISPR_CUSTOM) was used to select guide RNAs with relatively high activity and located close to the start codon or the stop codon of the target gene for N-terminal and C-terminal TAG-IN, respectively. In principle, other previously used co-injection markers for CRISPR, such as pSEM232 *(Pmlc-1::2xNLS::tagRFP::tbb-2 3’UTR)* or pRF4 (*rol-6(su1006)*) [28,55], can be used in place of the *dpy-10* co-conversion marker to enrich for F_1_ progeny carrying seed alleles. Annotated DNA sequences of the three small landing pads to generate C-terminal, N-terminal, or internal seed alleles are included in the S1 File. A list of guide RNAs, repair templates, and genotyping primers for seed alleles generated in this study is listed in S4 Table.

### Generation of tag donor strains

Tag donor strains, each of which carries a tag donor plasmid containing the tag as well as the SEC marker in an extrachromosomal array, were generated by microinjecting a mixture of DNA plasmids into the strain GE24 *pha-1*(*e2123ts*) hermaphrodites (viable at 15°C but not at room temperature). In general, a typical injection mixture contains a tag donor plasmid (25 ng/µL, unless specified), two HisCl1 negative selection markers (*myo-2p::HisCl1::GFP(or mScarlet-I)::tbb-2 3’UTR*, 2.5 ng/µl and *snt-1p::HisCl1-GFP(or mScarlet-I)::tbb-2 3’UTR*, 5 ng/µL), a *pha-1*(+) rescue plasmid (50 ng/µL), and DNA filler pBlueScript KS(+) for a total DNA concentration of 100 ng/µL. Injected P_0_ animals were transferred to room temperature to screen for surviving Rol animals in the next few generations to establish heritable extrachromosomal arrays as tag donor strains. Due to the SEC marker in the tag donor plasmid, the tag donor arrays exhibited a Rol phenotype and were resistant to hygromycin. For the convenience of maintaining the large library of tag donor arrays generated in this study, the *pha-1(+)* rescue plasmid was used as a co-injection marker for the donor array in the temperature-sensitive lethal *pha-1(e2123ts)* background. However, it is worth noting that new tag donor arrays can also be created in virtually any genetic background and maintained either by picking Rol animals or by applying hygromycin selection.

### Generation of an alternative transgene to express Flp recombinase in the germline

We used MosSCI to generate a single-copy germline-expressing Flp transgene, *tanSi334*, by injecting the plasmid pHW2134 into the strain EG6699. This inserted the DNA fragment *mex-5p::Flp(D5)::SL2::mNG::glh-2 3’UTR*, the same one used in the transgene *bqSi711*, into the *ttTi5605* locus on Chr II [25]. The *tanSi334* transgene was outcrossed six times with N2 to remove the background *unc-119(ed3)* mutation. Due to linkage, *bqSi711 IV* is not compatible with seed alleles on Chr IV for the TAG-IN workflow using genetic crossing (Fig 4A). *tanSi334* II serves as an alternative to *bqsi711* for those seed alleles on Chr IV and can also be used with seed alleles on chromosomes other than Chr II.

### Tag swapping using RMCE via microinjection

We followed a previously described injection protocol for RMCE [33]. Briefly, a tag donor plasmid containing the tag and the SEC cassette was diluted to 50 ng/μL in 10 μL nuclease-free water (NEB, Ipswich, MA) and then injected into double homozygous hermaphrodites carrying a seed allele of the target gene and *bqSi711*(a germline-expressing Flp transgene). Two injected P_0_ animals were plated per 60 mm NGM plate and cultured at room temperature. 400 μL of 5 mg/mL hygromycin B (GoldBio, St. Louis, MO) was added to each plate three days later to select for successful RMCE progenies. After another three days, Rol animals were then singled out onto new NGM plates without hygromycin until homozygous Rol animals were obtained in the next couple of generations. The SEC marker was excised by heat-shocking L1/L2 larvae at 30 °C for 18 hours [14]. Five days later, non-Rol progenies were singled out to establish final TAG-IN alleles.

### Tag swapping through RMCE via genetic crossing

N2 males were crossed with double-homozygous hermaphrodites carrying both a seed allele of the gene of interest and *bqSi711* (a germline-expressing Flp transgene); alternatively, *bqSi711* males were crossed with homozygous hermaphrodites carrying a seed allele of the gene of interest. Three days later (day 0 in Fig 4A), young males from the cross plates were picked out and crossed with a tag donor strain of choice. Typically, six plates were set up to ensure we got at least two independent TAG-IN alleles. The cross plates were stored at room temperature. Three days later, 400 μL of 5 mg/mL hygromycin B (GoldBio, St. Louis, MO) was added to each cross plate. This step selects for Rol animals, which were either cross progenies or self-reproduced progenies carrying the tag donor array. After another three days, 500 μL of 500 mM histamine dihydrochloride (HisCl, from GoldBio, St. Louis, MO) was added to each plate to select for positive plates containing successful TAG-IN lines. To support the growth of TAG-IN lines during histamine selection, it is important to ensure that each plate still has food before adding histamine. In the presence of histamine, the two negative selection markers, P*myo-2*::HisCl1 and P*snt-1*::HisCl1, robustly inhibit both pharyngeal muscle and the nervous system and thus render animals carrying the donor array unable to feed (pharyngeal pumping) or move normally. As a result, only animals with the tag as well as the SEC cassette inserted at the RMCE landing pad but without the donor array would survive and exhibit the Rol and robust pumping phenotypes. In our experience, such surviving Rol animals can appear as early as the day of histamine selection, and as late as a week after the selection, depending on the timing of the RMCE event. Typically, three days after histamine selection, 1-4 surviving Rol progenies per positive plate, which are candidate TAG-IN lines, were then singled out on individual NGM plates that did not contain hygromycin or histamine. The number of surviving Rol worms per positive plate to single out can be adjusted based on the number of positive plates and the numbers of independent lines to obtain. In the next generation, Rol animals were singled out again from candidate TAG-IN lines that produced ∼75% Rol progeny to obtain homozygous worms. The SEC was excised by heat-shocking L1/L2 larvae at 30°C for 18 hours [14]. Five days later, non-Rol progenies were singled out to establish final TAG-IN alleles. Note that TAG-IN alleles obtained from the same cross plate are considered to originate from the same line. During the initial single-out step following histamine selection, it can be advantageous (though not required) to select Rol animals that also contain the germline-expressing Flp transgene (indicated by green fluorescence in the germline due to mNG expression from the transgene), as expressing Flp recombinase in the germline for one additional generation may be helpful for reducing false positives by ensuring complete RMCE and full breakdown of the donor array [33]. All the tag donor strains generated in this study carry the temperature-sensitive lethal *pha-1(e2123ts)* mutation. As a result, final TAG-IN strains obtained from RMCE crosses would be either homozygous for *pha-1(+)* or heterozygous for *pha-1(ts)*. If this is a concern, additional outcrossing or genotyping may be needed to ensure complete loss of the *pha-1(ts)* mutation.

To confirm TAG-IN alleles were correct, genotyping PCR was performed using two outer primers specific to each tagging site to amplify the genomic region containing the FRT site, the tag, and the reverse FRT3 site as well as several hundred base pairs of upstream and downstream flanking sequences, the PCR products were analyzed using DNA gel electrophoresis and also further sequenced using Oxford Nanopore long-read sequencing (Plasmidsaurus, Eugene, OR).

### Nomenclature for TAG-IN alleles

According to *Caenorhabditis* nomenclature guidelines, each allele generated by genome editing at an endogenous locus, including seed alleles by CRISPR and TAG-IN alleles by RMCE, should be designated new allele name (*tan#* for our lab). Because each TAG-IN allele was generated by RMCE on a landing pad allele in a parental strain to insert a tag, TAG-IN alleles generated in this study were named as *gene(tan#1[description of the TAG-IN allele], *tan#2)*, where *tan#1* is the new TAG-IN allele and *tan#2* is the parental seed allele. Examples for TAG-IN nomenclature are as follows: for C-terminal (CT) tagging, *ceh-14(tan252[ceh-14::GFP CT TAG-IN], *tan236)*; for N-terminal (NT) tagging, *ceh-14(tan303[cGAL::SL2::ceh-14 NT TAG-IN], *tan259); for internal tagging (IT), ceh-14(tan386[ceh-14::mCherry IT TAG-IN], *tan386).* For brevity, the parental seed allele (*\*tan#2*) is often omitted when TAG-IN alleles are described in the main text. For example, *ceh-14(tan252[ceh-14::GFP CT TAG-IN])* is formally *ceh-14(tan252[ceh-14::GFP CT TAG-IN], *tan236)*, in which the CT seed allele *tan236* was used as the landing pad for RMCE to generate *tan252*.

### Behavioral assay

All the strains for the sleep assay carried a transgene (*syIs231*) that uses a heat-inducible promoter to conditionally overexpress LIN-3C/EGF upon heat shock [56]. Overexpression of EGF robustly induces sleep in *C. elegans*, during which animals show feeding and locomotion quiescence [57]. About 16 hours before the start of the assay, 25-30 L4 larvae were picked into a new NGM plate seeded with food. The next day, plates were sealed using Parafilm and then placed in a 33°C water bath for 15 minutes. After heat shock, the plates were immediately transferred to room temperature on a paper towel to dry the outside surface. Parafilm was removed, and the lid was half taken off to cool down the plates. The plates were not touched for the next 2 hours until they were ready to be scored. Lids of the plates were gently put on before scoring the plates. Sleep was scored on worms within the bacterial food patch by observing pharyngeal pumping right before and at 2 hours post-heat shock. Pumping quiescence was scored as no movement of pharyngeal grinder during 4 seconds of direct observation through a dissecting microscope. The observer was blinded to the genotypes of the worms.

### Microscopy

L4 animals were paralyzed in 20 μL of 30 mg/mL 2,3-Butanedione monoxime (BDM) solution (Sigma-Aldrich, St. Louis, MO) on a coverslip (VWR micro cover glasses, 22×22mm, No. 1.5). Extra BDM was used to wash out food in the drop of BDM containing worms. After cleaning up the food in the drop BDM, around 10 μL of BDM was left on the coverslip. To create imaging slides after most of the worms are immobilized, a freshly prepared 2% agarose pad on a glass slide (Gold Seal, 25 x 75 mm, 1 mm thick) was inverted onto the immobilized worms on the cover slides. All images were oriented with the anterior side of the worm to the left and the ventral side at the bottom. DIC and fluorescent images are taken using a Nikon Ti2-E Eclipse inverted microscope connected to a Prime 95B 25mm sCMOS camera and a SPECTRA X LED light engine (Lumencor, Beaverton, OR). Green and red fluorescent images were taken with standard GFP and mCherry filter cubes, respectively.

## Data availability

Many strains and plasmids generated in this paper have been deposited at the *Caenorhabditis* Genetics Center (CGC, https://cgc.umn.edu/) and Addgene (https://www.addgene.org/), respectively; others are available upon request.

## Acknowledgments

We thank the members of the Wang lab for comments on the manuscript, M. Nonet, P. Askjaer, J. Nance, and J. Ward for sharing plasmids and strains. We thank E. Jorgensen for sharing unpublished results. Some strains were provided by the CGC, which is funded by NIH Office of Research Infrastructure Programs (P40 OD010440). We thank WormBase for providing information about the *C. elegans* genome sequence and annotations. This work was supported by NIH NIGMS (R35GM150658), Whitehall Foundation, University of Wisconsin-Madison, and the Wisconsin Alumni Research Foundation. Some figures were created in https://BioRender.com.

## Supporting information

**S1 Fig. Schematics, DNA sequences, and protein products of C-terminal seed and TAG-IN alleles.**

**S2 Fig. Schematics, DNA sequences, and protein products of N-terminal seed and TAG-IN alleles**.

**S3 Fig. Schematics, DNA sequences, and protein products of internal seed and TAG-IN alleles**

**S4 Fig. Table showing the success rates of generating TAG-IN alleles through RMCE by injection.**

**S5 Fig. Expression of *ceh-14* N-terminal co-expression TAG-IN allele in the ALA neuron at the L3 stage.**

**S6 Fig. Functional validation of a new germline-expressing Flp transgene and evaluation of iterative RMCE efficiency using TAG-IN alleles as landing pads.**

**S1 Table. RMCE efficiency by genetic crossing.**

**S2 Table. Strains used in this study.**

**S3 Table. Plasmids used in this study.**

**S4 Table. Oligos used in this study.**

**S1 File. Annotated DNA sequences of seed allele landing pads, TAG-IN donor plasmids, and TAG-IN destination vectors.**

## Notes

https://www.addgene.org/browse/article/28274025/

### Competing Interest Statement

The authors have declared no competing interest.

### Summary of Updates

title and main text updated; TAG-IN at multiple new loci tested; supplemental files updated.

