## Supplementary Figures S1-S6 for "TAG-IN, a modular strategy for swappable endogenous gene tagging in *Caenorhabditis elegans*"

**Schematic of a C-terminal seed allele:**
the target gene, immediately upstream of its stop codon-(GCA G)-SD-FRT-(mini splice acceptor)-(GA TCC)-rFRT3-its native stop codon and 3’ UTR

**DNA sequence of a C-terminal seed allele:**
the target gene-GCAGgtaagtttaaacatgattttactaacgaagttcctattctctagaaagtataggaacttcttt
tcagGATCCGAAGTTCCTATACTATTTGAAGAATAGgaacttc-its native stop codon and 3’UTR

**Protein product**: POI-linker(“AGSEVPILFEE”)

**Schematic of a C-terminal fusion TAG-IN allele:**
the target gene, immediately upstream of its stop codon--(GCA G)-SD-FRT-SA-(GA TCC GGT GGC AGT GGA GGT ACC GGC GGA AGC GGT, flexible linker1)-(ATG)-sequence of the tag-(AAG)-SD’-lox site-SA’-(GGT)-flexible linker2-(ACG)-STOP-rFRT3-its native stop codon and 3’UTR

**DNA sequence of a C-terminal fusion TAG-IN allele (use loxP as an example):**
the target gene-GCAGgtaagtttaaacatgattttactaacgaagttcctattctctagaaagtataggaacttctaa
ctaatctgatttaaattttcagGATCCGGTGGCAGTGGAGGTACCGGCGGAAGCGGTATGsequence of the tagAAGg
taagtttaaaataacttcgtatagcatacattatacgaagttattttcagGGTGGCAGCGGAGGTACCGGCGGTAGTGGAGGC
ACGTAAgaagttcctatactatttgaagaataggaacttc-its native stop codon and 3’UTR

**Protein product**: POI-flexible linker1(“AGSGGSGGTGGSG”)-tag-flexible linker2#(“KGGSGGTGGSGGT”), as a single fusion protein

**Schematic of a C-terminal co-expression TAG-IN allele:**
the target gene, immediately upstream of its stop codon--(GCA G)-SD-FRT-SA-(GA TCC GGT GGC AGT GGA GGT ACC GGC GGA AGC GGT, flexible linker1)-STOP-SL2-Kozak sequence(aaaa)-(ATG)-sequence of the tag-(AAG)-SD’-lox site-SA’-(GGT)-flexible linker2-STOP-rFRT3-its native stop codon and 3’UTR

**DNA sequence of a C-terminal co-expression TAG-IN allele (use lox511I as an example):**
the target gene-GCAGgtaagtttaaacatgattttactaacgaagttcctattctctagaaagtataggaacttctaa
ctaatctgatttaaattttcagGATCCGGTGGCAGTGGAGGTACCGGCGGAAGCGGTTAAgctgtctcatcctactttcacct
agttaactgcttgtcttaaaatctatgcttctctttagtatctaaaattttcctagaagcttacaagtatataaatggtctct
tctcaataaaggttgtatatttattcatcttattgaatctgccatttcctcgtttttgcgagtttatataccttccaattttc
tttctattgtattttcaacttctaattttaattcagggaaactgcttcaacgcatcaaaaATGsequence of the tagA
AGgtaagtttaaaataacttcgtataatgtatactatacgaagttattttcagGGTGGCAGCGGAGGTACCGGCGGTAGTGGA
GGCACGTAAgaagttcctatactatttgaagaataggaacttc-its native stop codon and 3’UTR

**Protein products**: POI-flexible linker1(“AGSGGSGGTGGSG”) and tag-flexible linker2#(“KGGSGGTGGSGGT”), as two separate proteins that are co-expressed in the same cells

**S1 Fig. Schematics, DNA sequences, and protein products of C-terminal seed and TAG-IN alleles**. Annotations and DNA sequences are highlighted using a matching color scheme. The “sequence of the tag” denotes the DNA sequence encoding the tag, excluding both the start codon and the stop codon. #, if a stop codon is included immediately downstream of the tag in the donor plasmid, no flexible linker2 is present at the C-terminus of the tag (see an example in Fig 2C). DNA sequences in uppercase indicate coding regions and are spaced to reflect the reading frame, while sequences expected to be spliced out of the mature mRNAs are underlined. The “SD’-lox site-SA’” section refers to the 51-bp residual intron left after self-removal of the SEC marker via heat shock. SD and SD’, splice donor; SA and SA’, splice acceptor; FRT, Flp Recognition Target (34 bp); rFRT3, reverse-oriented FRT variant 3; lox site, Cre recombination site (34 bp); SL2, SL2 intercistronic sequence from the *gpd-2/gpd-3* operon; POI, protein of interest. Quoted sequences within parentheses are the actual peptide sequences of the linkers in the final protein products. Annotated DNA sequences of the 110-bp C-terminal landing pad and all C-terminal tag donor plasmids are included in Supplementary S1 File.

**Schematic of an N-terminal seed allele:**
the target gene, immediately upstream of its start codon-FRT-(GGA G)-(mini splice donor)-rFRT3-SA-(GC ACG)-the native start codon

**DNA sequence of an N-terminal seed allele:**
the target gene’s 5’UTR-gaagttcctattctctagaaagtataggaacttcGGAGgtaagtttgaagttcctatact
atttgaagaataggaacttctaactaatctgatttaattttcagGCACG-its native start codon

**Protein product**: POI

**Schematic of an N-terminal fusion TAG-IN allele:**
the target gene, immediately upstream of the start codon-FRT-(AAG)-SD’-lox site-SA’-(GGT)-(ATG)-sequence of the tag-(ACG)-(GGC AGC GGA GGT ACC GGC GGT AGT GGA G, flexible linker)-SD-rFRT3-SA-(GC AGG)-its native start codon

**DNA sequence of an N-terminal fusion TAG-IN allele (use loxP as an example):**
the target gene’s 5’UTR-gaagttcctattctctagaaagtataggaacttcaaggtaagtttaaaataacttcgta
tagcatacattatacgaagttattttcagggtATGsequence of the tagACGGGCAGCGGAGGTACCGGCGGTAGTGG
AGgtaagtttaaacatgattttactaacgaagttcctatactatttgaagaataggaacttctaactaatctgatttaaattt
tcagGCACG-its native start codon

**Protein product**: tag-flexible linker(“TGSGGTGGSGGT”)-POI, as a single fusion protein

**Schematic of an N-terminal co-expression TAG-IN allele:**
the target gene, immediately upstream of its start codon-FRT-(AAG)-SD’-lox site-SA’-(GGT)-(ATG)-sequence of the tag-(ACG)-STOP-SL2-Kozak sequence(aaaa)-(ATG GGC AGC GGA GGT ACC GGC GGT AGT GGA G, flexible linker)-SD-rFRT3-SA-(GC ACG)-its native start codon of the target gene

**DNA sequence of an N-terminal co-expression TAG-IN allele (use lox511I as an example):**
the target gene’s 5’UTR-gaagttcctattctctagaaagtataggaacttcaaggtaagtttaaaataacttcgta
tagcatacattatacgaagttattttcagggtATGsequence of the tagACGTAAgctgtctcatcctactttcacct
agttaactgcttgtcttaaaatctatgcttctctttagtatctaaaattttcctagaagcttacaagtatataaatggtctct
tctcaataaaggttgtatatttattcatcttattgaatctgccatttcctcgtttttgcgagtttatataccttccaattttc
tttctattgtattttcaacttctaattttaattcagggaaactgcttcaacgcatcaaaaATGGGCAGCGGAGGTACCGGCGG
TAGTGGAGgtaagtttaaacatgattttactaacgaagttcctatactatttgaagaataggaacttctaactaatctgattt
aaattttcagGCACG-its native start codon

**Protein products:** tag-(“T”) and flexible linker(“MGSGGTGGSGGT”)-POI, as two separate proteins that are co-expressed in the same cells

**S2 Fig. Schematics, DNA sequences, and protein products of N-terminal seed and TAG-IN alleles.** Annotations and DNA sequences are highlighted using a matching color scheme. The “sequence of the tag” denotes the DNA sequence encoding the tag, excluding both the start codon and the stop codon. DNA sequences in uppercase indicate coding regions and are spaced to reflect the reading frame, while sequences expected to be spliced out of the mature mRNAs are underlined. The “SD’-lox site-SA’” section refers to the 51-bp residual intron left after self-removal of the SEC marker via heat shock. SD and SD’, splice donor; SA and SA’, splice acceptor; FRT, Flp Recognition Target (34 bp); rFRT3, reverse-oriented FRT variant 3; lox site, Cre recombination site (34 bp); SL2, SL2 intercistronic sequence from the *gpd-2/gpd-3* operon; POI, protein of interest. Quoted sequences within parentheses are the actual peptide sequences of the linkers in the final protein products. Annotated DNA sequences of the 110-bp N-terminal landing pad and all N-terminal tag donor plasmids are included in Supplementary S1 File.

**Schematic of an internal seed allele:**
the target gene, 5’ exon-(GG)-FRT-(GGA G)-(mini splice donor)-rFRT3-SA-(GC ACG)-the target gene, 3’ exon

**DNA sequence of an internal seed allele (in-frame insertion into an exon):**
the target gene, 5’ exon-GGGAAGTTCCTATTCTCTAGAAAGTATAGGAACTTCGGAGgtaagtttgaagttcctatactatttgaagaataggaacttctaactaatctgatttaattttcagGCACG-the target gene, 3’ exon

**Protein product**: (POI-N)-linker(“GKFLFSRKYRNFGGT”)-(POI-C)

**Schematic of an internal fusion TAG-IN allele:**
the target gene, 5’ exon-(GG)-FRT-(AAG)-SD’-lox site-SA’-(GGT)-(ATG)-sequence of the tag-(ACG)-(GGC AGC GGA GGT ACC GGC GGT AGT GGA G, flexible linker)-SD-rFRT3-SA-(GC AGG)-the target gene, 3’ exon

**DNA sequence of an internal fusion TAG-IN allele (use loxP as an example):**
the target gene, 5’ exon-GGGAAGTTCCTATTCTCTAGAAAGTATAGGAACTTCAAGgtaagtttaaaataacttcgtatagcatacattatacgaagttattttcagGGTATGsequence of the tagACGGGCAGCGGAGGTACCGGCGGTAGTGGAGgtaagtttaaacatgattttactaacgaagttcctatactatttgaagaataggaacttctaactaatctgatttaaattttcagGCACG-the target gene, 3’ exon

**Protein product**: (POI-N)-linker(“GKFLFSRKYRNFKG”)-tag-flexible linker(“TGSGGTGGSGGT”)-(POI-C), as a single fusion protein

**S3 Fig. Schematics, DNA sequences, and protein products of internal seed and TAG-IN alleles.** Annotations and DNA sequences are highlighted using a matching color scheme. Note that the 112-bp internal landing pad must be inserted in-frame into an exon of the target gene to generate an internal seed allele, in which the exon is split into 5’ and 3’ exons. The “sequence of the tag” denotes the DNA sequence encoding the tag, excluding both the start codon and the stop codon. DNA sequences in uppercase indicate coding regions and are spaced to reflect the reading frame, while sequences expected to be spliced out of the mature mRNAs are underlined. The 12-aa segment (“GKFLFSRKYRNF”) of the linker is derived from the 36-bp recombination scar DNA sequence (GG + 34-bp FRT site, wave underlined). The “SD’-lox site-SA’” section refers to the 51-bp residual intron left after self-removal of the SEC marker via heat shock. SD and SD’, splice donor; SA and SA’, splice acceptor; FRT, Flp Recognition Target (34 bp); rFRT3, reverse-oriented FRT variant 3; lox site, Cre recombination site (34 bp); SL2, SL2 intercistronic sequence from the *gpd-2/gpd-3* operon; POI, protein of interest. Quoted sequences within parentheses are the actual peptide sequences of the linkers in the final protein products. An annotated DNA sequence of the 112-bp internal landing pad is included in Supplementary S1 File.

| Strains for injection | Tag | Tag donor plasmid | TAG-IN success rate (positive plates/total injection plates) |
|---|---|---|---|
| *bqSi711 IV; ceh-14(tan236[CT seed allele]) X* | SL2::cGAL | pHW2023 | 33% (3/9) |
| | GFP | pHW2024 | 10% (1/10) |
| | mCherry | pHW2025 | 27% (3/11) |
| *bqSi711 IV; ceh-14(tan259[NT seed allele]) X* | cGAL::SL2 | pHW2115 | 58% (7/12) |
| | GFP | pHW2119 | 60% (3/5) |
| Average success rate | | | 36% (17/47) |

**S4 Fig. Table showing the success rates of generating TAG-IN alleles through RMCE by injection.** Individual tag donor plasmids were injected into *ceh-14* seed alleles carrying the Flp-expressing transgene *bqSi711*. Two injected $P_0$ animals were placed on each injection plate.

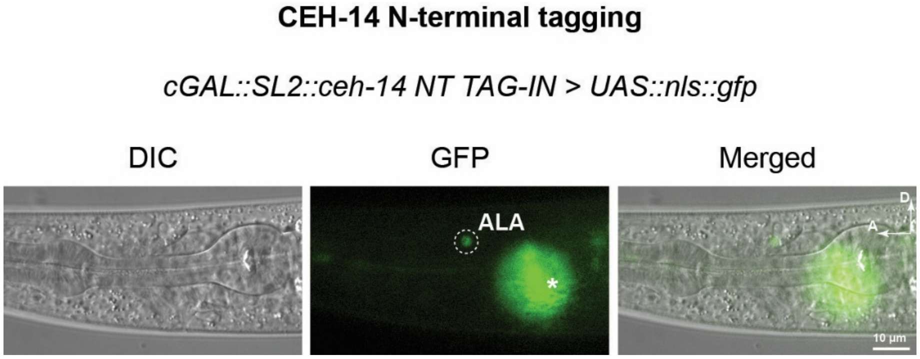


**S5 Fig. Expression of *ceh-14* N-terminal co-expression TAG-IN allele in the ALA neuron at the L3 stage.** DIC, Fluorescent, and merged images of the head region of the strain *cGAL::SL2::ceh-14 NT TAG-IN; UAS::nls::gfp* are shown. cGAL binds to UAS to drive expression of *nls::gfp* in the *ceh-14*-expressing cells. The white circle indicates the location of the ALA neuron. The asterisk (*) marks an out-of-focus neuron that is much brighter than the ALA neuron. Images were taken from animals at the L3 stage. A, anterior; D, dorsal. Scale bar: 10 μm.

**A**

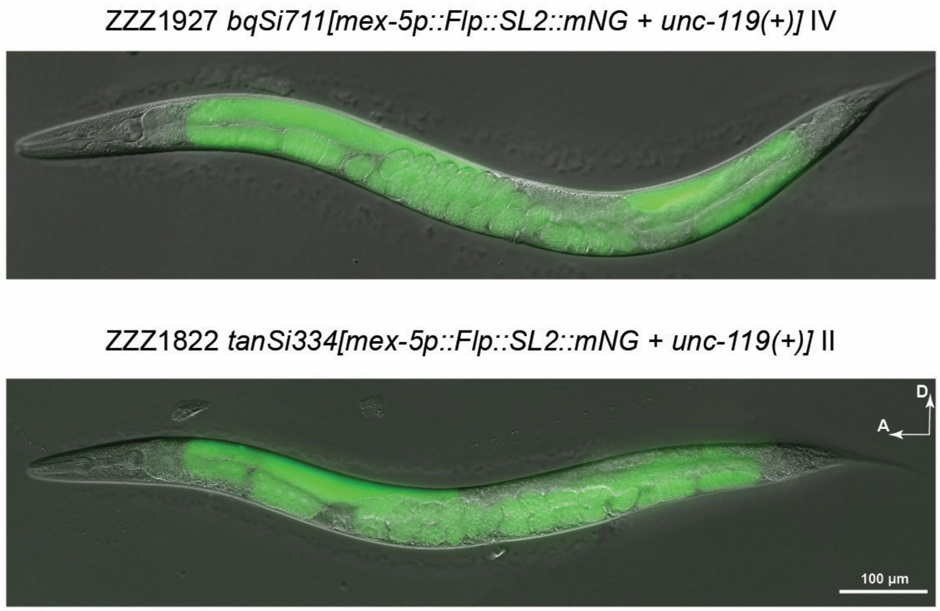


**B**

| Flp transgene | TAG-IN allele as seed allele | Tag donor strains | TAG-IN success rate (positive plates / total cross plates) |
|---|---|---|---|
| *bqSi711 IV* | *ceh-14(tan307[gfp::ceh-14* NT TAG-IN]*) X* | ZZZ1779 (mScarlet-I) | 58% (7/12) |
| *bqSi711 IV* | | ZZZ1776 (mCherry) | 50% (6/12) |
| *tanSi334 II* | | ZZZ1779 (mScarlet-I) | 42% (5/12) |

**S6 Fig. Functional validation of a new germline-expressing Flp transgene and evaluation of iterative RMCE efficiency using TAG-IN alleles as landing pads.** **(A)** Images showing mNG expression in the germline of adults carrying the indicated single-copy Flp transgenes on different chromosomes. A, anterior; D, dorsal. Scale bar, 100 µm. **(B)** Table showing the success rates of iterative RMCE used to generate new TAG-IN alleles through genetic crossing of an existing *gfp::ceh-14* N-terminal TAG-IN allele (serving as a landing pad, like the original *ceh-14* N-terminal seed allele) with the indicated Flp-expressing transgenes and tag donor strains.
